# Signal Strength Aware Latent Spaces Reveal Molecularly Distinct Substructures within Human Kidney Tissue

**DOI:** 10.64898/2026.01.15.699686

**Authors:** Paul-Louis Delacour, Lukasz G. Migas, Melissa A. Farrow, Haichun Yang, Agnes B. Fogo, Jeffrey M. Spraggins, Raf Van de Plas

## Abstract

As datasets grow increasingly high-dimensional and complex, distinguishing a condensed set of interpretable underlying factors becomes essential. In spatial omics, for example, hundreds to thousands of molecular features per observation promise unprecedented biological insight. However, without meaningful latent representations, that potential remains markedly untapped. We propose a new approach based on the beta-variational autoencoder and kernel density estimation to dissect data along independent, uncertainty-aware, and interpretable (yet non-linear) latent axes. We include a novel comparative-latent-traversal algorithm to translate latent findings back into the original measurement context. Demonstrating on imaging mass spectrometry-based molecular imaging of human kidney, the approach’s disentangling properties are shown to impress a latent space structure that separates signal strength from relative signal content, offering exceptional chemical insight. Our approach uncovers unexpected subdivisions within kidney proximal tubules, confirmed to be biological, and reveals hereto-unknown lipid species differentiating them. This confirms our workflow’s potential as an interpretation-and-hypothesis-generating discovery tool.

## 1 Introduction

Spatial biology technologies are advancing rapidly, enabling deeper molecular coverage with ever-greater resolution and yielding data of unprecedented molecular and spatial specificity. By capturing molecular relationships down to cellular resolution, these large multidimensional datasets offer deep insights into biological systems. However, their complexity makes deciphering underlying patterns challenging, particularly considering nonlinear relationships. To leverage the full potential of these intricate datasets, new methods are needed to discover and, more importantly, interpret the relationships underlying the measurements. Placing such findings in the original measurement context is essential to understanding the driving biology behind them. Here, we introduce a workflow specifically designed to uncover a generative mechanism underlying the data, thereby facilitating human interpretation. We demonstrate the effectiveness of this approach in molecular imaging, a field characterized by complex, high-dimensional data. Specifically, we apply our method to imaging mass spectrometry (IMS) [1,2], a highly multiplexed modality, used for untargeted molecular mapping of human tissue. As a spatial omics technology, IMS acquires mass spectra pixel-by-pixel across a sample surface. A single experiment maps the tissue-wide abundance of hundreds to thousands of endogenous molecules (*e.g*., metabolites, lipids, glycans, proteins) simultaneously, without requiring prior chemical labeling. The ability of our approach to reveal nuanced structure within IMS data can help drive discovery of biomolecular pathways active in disease, reveal patient-specific insights for precision medicine, and offer a novel automated means of hypothesis-generation for biomedical research.

Dimensionality reduction methods have become the norm for overcoming the complexity of multidimensional data, mapping the variation of high-dimensional measurements into a lower-dimensional latent space. Capturing underlying patterns in data is critical to many areas, from satellite-based remote sensing [3] and industrial process monitoring [4] to transcriptomics [5] and climate science [6]. However, it is often difficult to link observations in the low-dimensional representation to specific features of the measurements. Nevertheless, the ability to map latent space features back into the original measurement space is essential to aiding interpretation. Linear methods such as principal component analysis (PCA) [7–10] and non-negative matrix factorization (NMF) [11,12] implicitly deliver such an approximate inverse mapping, facilitating direct interpretation. However, many relationships underlying high-dimensional molecular data are not necessarily linear and cannot be captured accurately by such relatively basic models. Therefore, nonlinear mappings into a latent space are needed, as with t-distributed stochastic neighbor embedding (t-SNE) [13] or uniform manifold approximation and projection (UMAP) [14,15]. Unfortunately, inverting from a nonlinear latent space to reveal corresponding variation in the measurement space is much less straightforward and sometimes impossible. In biomedical research, omics measurements using transcriptomics or mass spectrometry routinely report hundreds to thousands of molecular features per observation. While such deep characterizations promise unprecedented insight into biological processes and molecular phenotypes, part of that potential remains untapped unless meaningful information can be extracted from latent representations of the data. Specifically, latent space axes should ideally capture biologically relevant aspects, *e.g*., variation distinguishing cell types, cellular niches, or healthy-versus-diseased tissue, or report molecular abundance. It also requires a method that, after capturing nonlinear relationships in a latent form, allows them to be mapped back to the measurement space for biological understanding. One class of models suitable for achieving both goals is the autoencoder. Autoencoders, and more specifically variational autoencoders (VAEs) [16], simultaneously create forward (encoding) and backward (decoding) nonlinear mappings between the native and latent spaces, enabling insights from a dataset’s low-dimensional representation to be cast back into the context of the original measurements.

VAEs, in particular *β*-variational autoencoders (*β*-VAEs) [17], are generative methods for dimensionality reduction. Not only do they cast high-dimensional data into a lower-dimensional latent space, but a measurement is mapped to a distribution rather than a single point in that latent space, and the relationship between the latent space and native measurements is explicitly modeled (Figure 1a). Moreover, *β*-VAEs are designed to enforce statistical independence between the latent factors they extract. This compels the resulting latent space to evolve towards a structure in which each latent dimension corresponds to a different generative factor within the data. In a biological context, such independent latent dimensions can, for example, report distinctions between biological processes, molecular pathways, or cell types. The generative nature additionally captures uncertainty, which is crucial when the number of observations is small, as in most omics experiments. Specifically, we will show that, at least for IMS, the disentangling properties of the *β*-VAE tend to give the latent space a systematic structure that separates signal content from signal strength.

**Fig. 1:**
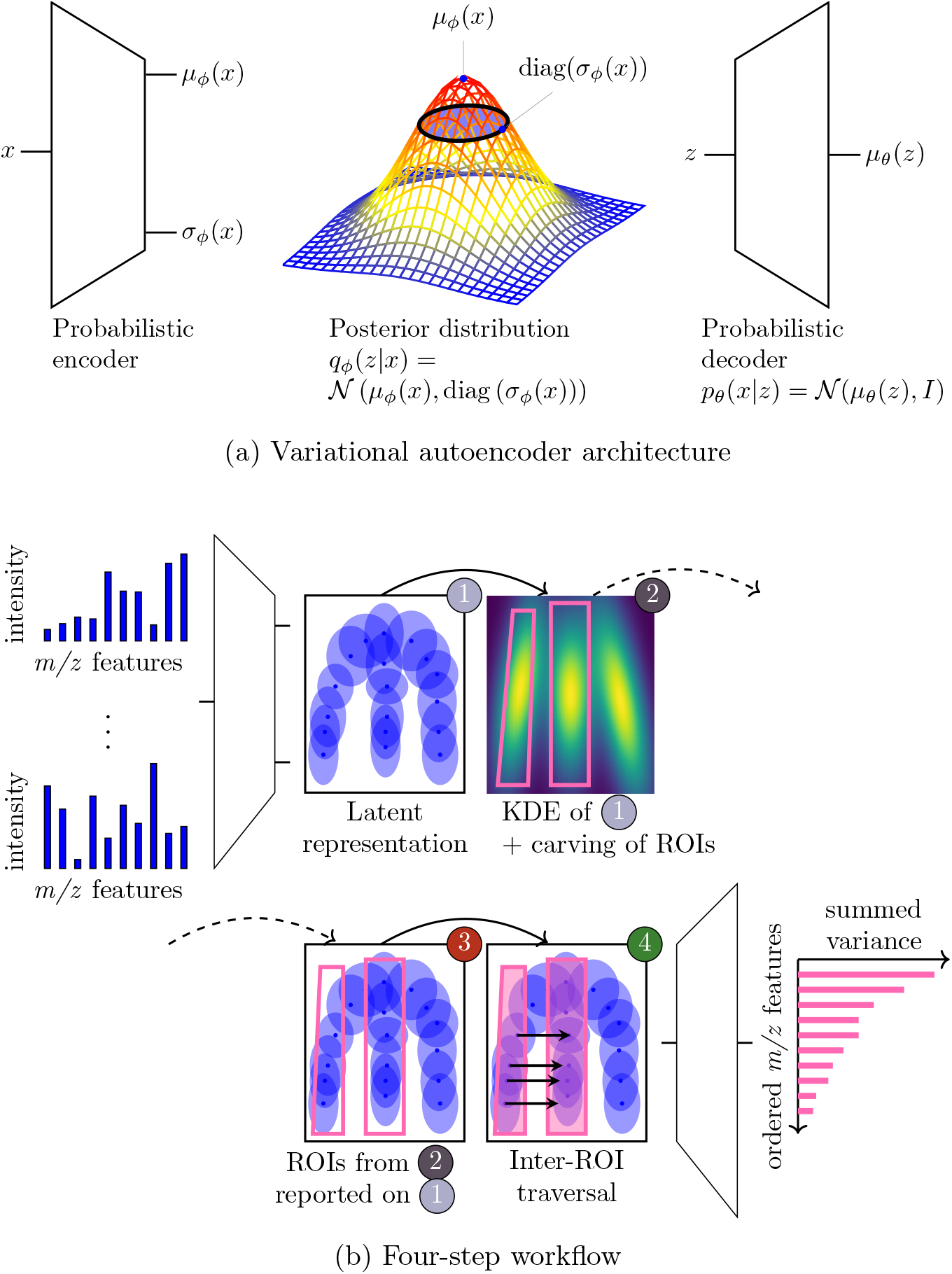
(1a) Architecture of traditional variational autoencoder (VAE). (1b) Overview of the workflow, as applied to an IMS-dataset. The workflow (details in section 4) delivers three outputs: (i) a point-cloud latent representation of the mass spectra in the dataset (with uncertainty captured as an accompanying standard deviation around each mean point); (ii) a rasterized latent representation; and (iii) an ordering of measurement space (*m/z*) features as a function of their accumulated variance when traversing from latent region-of-interest (ROI) to ROI. Result (i) captures probabilistic structure underlying the data using a given number of latent dimensions and disentangles generative factors, (ii) reports probabilistic density and facilitates recognition of specific latent ROIs, and (iii) identifies the most varying ion species between ROIs, *i.e*., the molecular species differentiating between the tissue structures corresponding to those ROIs.

Herein, we develop a *β*-VAE-based workflow that provides a latent space that not only captures global structure of complex, high-dimensional IMS data, but also offers independent, interpretable axes. We also introduce a new algorithm to cast latent findings back to the original measurement features, in this case for IMS. Although autoencoders [18,19] and VAEs [20,21] have been successfully applied to IMS, these approaches do not yield the independent axes we need for interpretation, motivating a necessary progression towards the *β*-VAE. Performance of our workflow is demonstrated using IMS measurements of human kidney acquired as part of the Human Bio-Molecular Atlas Program (HuBMAP), a large-scale, multi-institutional initiative generating foundational molecular atlases of human tissues at cellular resolution [22,23]. The ability of our method to reveal, in an interpretable latent space, previously unknown relationships between molecular measurements and biologically relevant tissue structures underscores its potential as an interpretation and hypothesis-generating discovery tool.

## 2 Results

Our workflow entails four stages (Figure 1b). First, a *β*-VAE is trained to learn a probabilistic latent space (Methods–step 1). Its variational aspects capture the intrinsic variability of the data. Each IMS pixel, *i.e*., mass spectrum, is cast into a low-dimensional space where it is represented by a mean location and one associated standard deviation around it (Figure 1a). Next, we introduce kernel density estimation (KDE) as a means to reveal structure in this probabilistic latent space and to make it human-perusable (Methods–step 2). The KDE representation enables users to recognize areas of high probabilistic density and to segment or ‘carve out’ specific latent regions-of-interest (ROIs) (Methods–step 3). A high-density ROI can be mapped by pixel membership to the measurement space’s spatial domain, often delineating key tissue features and cell types. However, since latent ROIs do not reveal which molecular species are responsible for differentiating their corresponding tissue structures, mapping latent variation onto the measurement space’s spectral domain is an essential next step. Our novel latent-traversal algorithm facilitates this by projecting latent differences onto measurement space features (Methods–step 4), effectively identifying which molecular species differentiate between user-specified ROIs. Methods section 4 provides details.

We apply our workflow to five distinct datasets (Table 1). In case study 1 (section 2.1), we demonstrate on a human kidney IMS dataset our method’s ability to disentangle underlying (biology-relevant) latent structures. In case study 2 (section 2.2), we use a synthetic dataset with a pre-specified spatial mixture of spectral signatures to examine how certain types of measurement variation manifest themselves in the latent space. The synthetic case study is also used to reinforce an observation from case study 1: for IMS-like measurements, our workflow tends to learn latent factors that separate relative from absolute variation, *i.e*., that separate signal content from signal strength or signal-to-noise-ratio (SNR). We will refer to such a space as a signal strength aware latent (SiSAL) space. This observation is consistent with the previously established spiked mixture model [24], a statistical model for IMS and hyperspectral imaging measurements that also captures variation in terms of SNR and signal content. After substantiating our workflow’s ability to capture genuine biological structure in its latent representation and its tendency towards imposing a SiSAL organization on that representation, we revisit case study 1’s kidney dataset (section 2.3). There, we demonstrate our algorithm for casting latent variation back into the original measurements’ feature space. Finally, we demonstrate the capacity of our method for hypothesis generation (section 2.4). Using lipid imaging data collected from multiple patients (case studies 3-5), we differentiate anatomical subdivisions within the kidney’s proximal tubules (PT) based on their SiSAL-revealed molecular profiles. These case studies conclude with our latent-traversal algorithm identifying a previously unknown lipid species driving the distinction between PT segments. The SiSAL Python package that implements our approach, along with case study 1 materials and the synthetic data experiment, is provided at https://github.com/vandeplaslab/sisal.

**Table 1:**
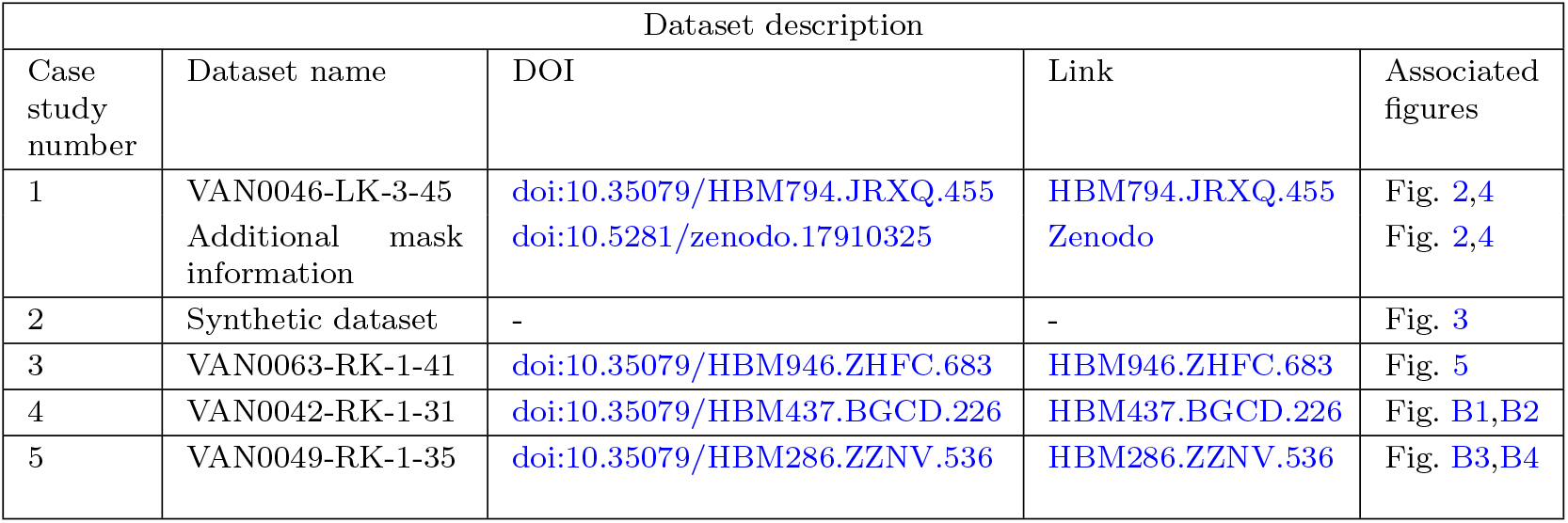
Overview of case studies and corresponding datasets. All human kidney datasets are part of the Human Bio-Molecular Atlas Program (HuBMAP) [22]. Additional information, such as FTU masks, is made available as a Zenodo repository. The SiSAL Python package is available at: https://github.com/vandeplaslab/sisal.

| Dataset description |  |  |  |  |
| --- | --- | --- | --- | --- |
| Case study number | Dataset name | DOI | Link | Associated figures |
| 1 | VAN0046-LK-3-45<br>Additional mask information | [doi:10.35079/HBM794.JRXQ.455](https://doi.org/10.35079/HBM794.JRXQ.455)<br>[doi:10.5281/zenodo.17910325](https://doi.org/10.5281/zenodo.17910325) | [HBM794.JRXQ.455](https://doi.org/10.35079/HBM794.JRXQ.455)<br>[Zenodo](https://zenodo.org/record/17910325) | Fig. 2,4<br>Fig. 2,4 |
| 2 | Synthetic dataset | - | - | Fig. 3 |
| 3 | VAN0063-RK-1-41 | [doi:10.35079/HBM946.ZHFC.683](https://doi.org/10.35079/HBM946.ZHFC.683) | [HBM946.ZHFC.683](https://doi.org/10.35079/HBM946.ZHFC.683) | Fig. 5 |
| 4 | VAN0042-RK-1-31 | [doi:10.35079/HBM437.BGCD.226](https://doi.org/10.35079/HBM437.BGCD.226) | [HBM437.BGCD.226](https://doi.org/10.35079/HBM437.BGCD.226) | Fig. B1,B2 |
| 5 | VAN0049-RK-1-35 | [doi:10.35079/HBM286.ZZNV.536](https://doi.org/10.35079/HBM286.ZZNV.536) | [HBM286.ZZNV.536](https://doi.org/10.35079/HBM286.ZZNV.536) | Fig. B3,B4 |

### 2.1 Mapping molecular measurements to a SiSAL space

In case study 1, we analyzed IMS data from human kidney tissue (HuBMAP VAN0046-LK-3-45), considering each pixel (*i.e*., preprocessed mass spectrum) as a separate data point and using 80% of the total dataset for *β*-VAE training. Figure 2 shows steps 1 through 3 (Figure 4 depicts step 4). The resulting latent space (step 1) achieved a *>*100-fold reduction in dimensionality (212-to-2). However, the probabilistic nature of its latent representation (depicted using standard-deviation ellipses in Figure 2a) made discerning latent structure nontrivial. When KDE was performed (step 2), several distinct vertical latent density striations became clear (Figure 2b). The density-revealing view offered by KDE allowed latent regions-of-interest (ROIs) to be ‘carved out’ (step 3). After delineation on the KDE representation (examples in Figure 2b), the ROIs were copied to the variational representation (Figure 2c). There, the mean latent locations of pixels and their presence inside ROIs were used to project each latent ROI to the spatial domain of the measurement space (Figure 2d). Each carving of the latent space resulted in a spatial segment that delineated a distinct kidney tissue structure. Figure 2e shows one of 212 ion images in this dataset, namely mass-to-charge-ratio (*m/z*) 556.306, an ion that localized to the glomeruli. The correlation between this glomerulus-reporting ion and the orange ROI segment (Figure 2d, see arrows) suggested that the orange latent ROI in Figure 2c captured (at least) data variation that aligns with signals that localize to the glomeruli. Considering the ROIs’ shapes, molecular variation that differentiates tissue structures appears to be captured along the *z*_1_-axis, while a pixel’s SNR seems to be encoded along the *z*_2_-axis (see ellipse sizes). This suggested presence of a SiSAL space, prompting a confirmatory examination in case study 2.

**Fig. 2:**
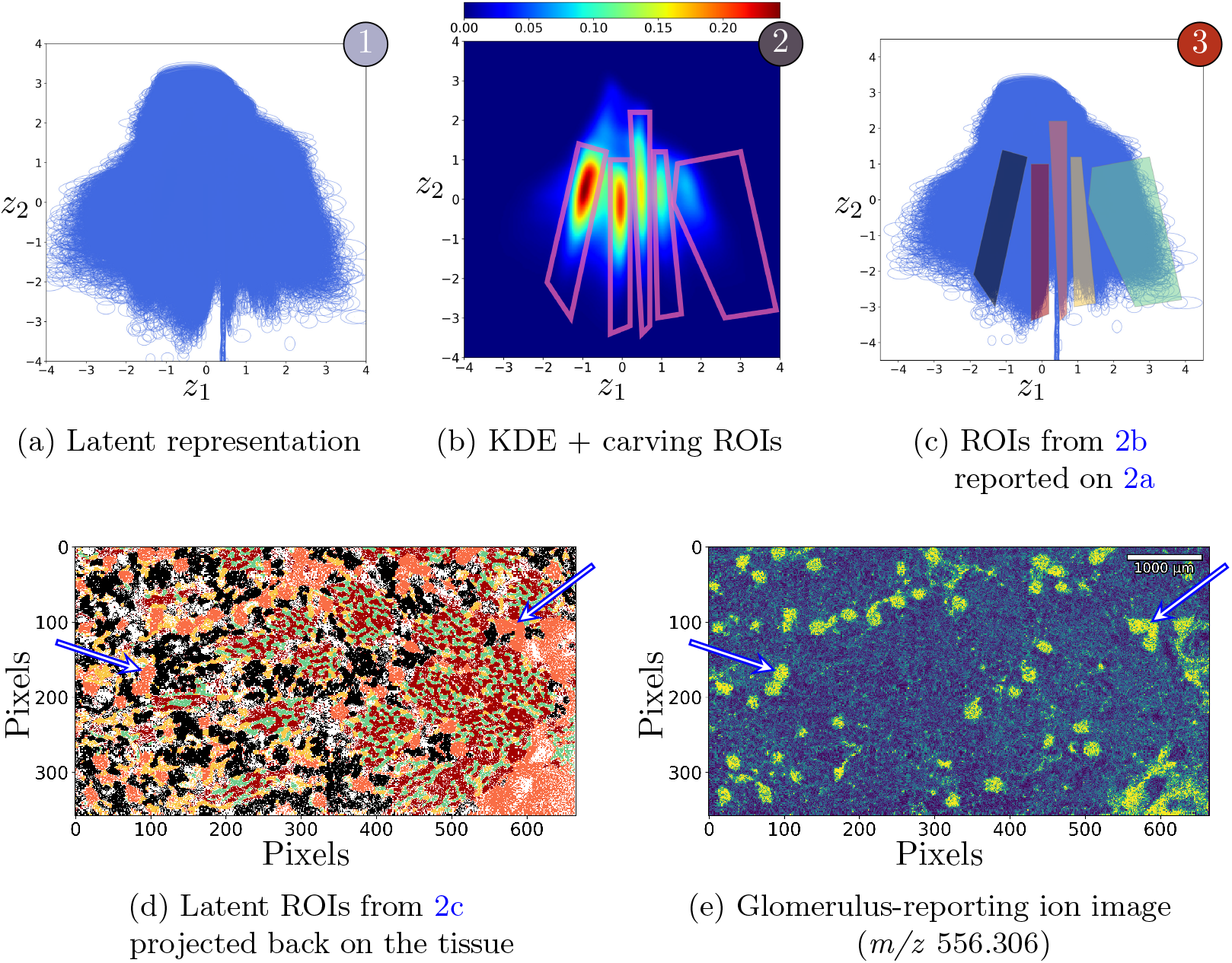
Workflow applied to case study 1’s human kidney IMS data. (2a) Post-step-1: the learned generative latent space. Each spectrum is mapped to a mean location and standard-deviation ellipse in a 2-D plane. (2b) Post-step-2: the density-based representation obtained through KDE and manually defined segmentations of example latent ROIs (polygons). (2c) Post-step-3: ROI segmentations from panel 2b copied to the latent representation of panel 2a, assigning different colors to each ROI-polygon. (2d) Projecting the latent ROIs back to the spatial domain of the measurement space shows each ROI corresponding to a different kidney tissue substructure. (2e) A glomerulusreporting ion feature, *m/z* 556.306, correlates to the orange ROI areas (see arrows), suggesting that the orange ROI-polygon in panel 2c captures (at least) glomerular variation.

### 2.2 Confirming the SiSAL nature of the developed latent space

To better understand how measurements (*e.g*., IMS spectra) lead to the observed latent structure, we generated a synthetic dataset from known underlying signatures. This dataset modeled a spatial domain, with different regions generating distinct qualitative spectra, and a pixel-specific quantitative signal strength. Figure 3a shows how this synthetic dataset was constructed. Section 4.3 provides a detailed description.

**Fig. 3:**
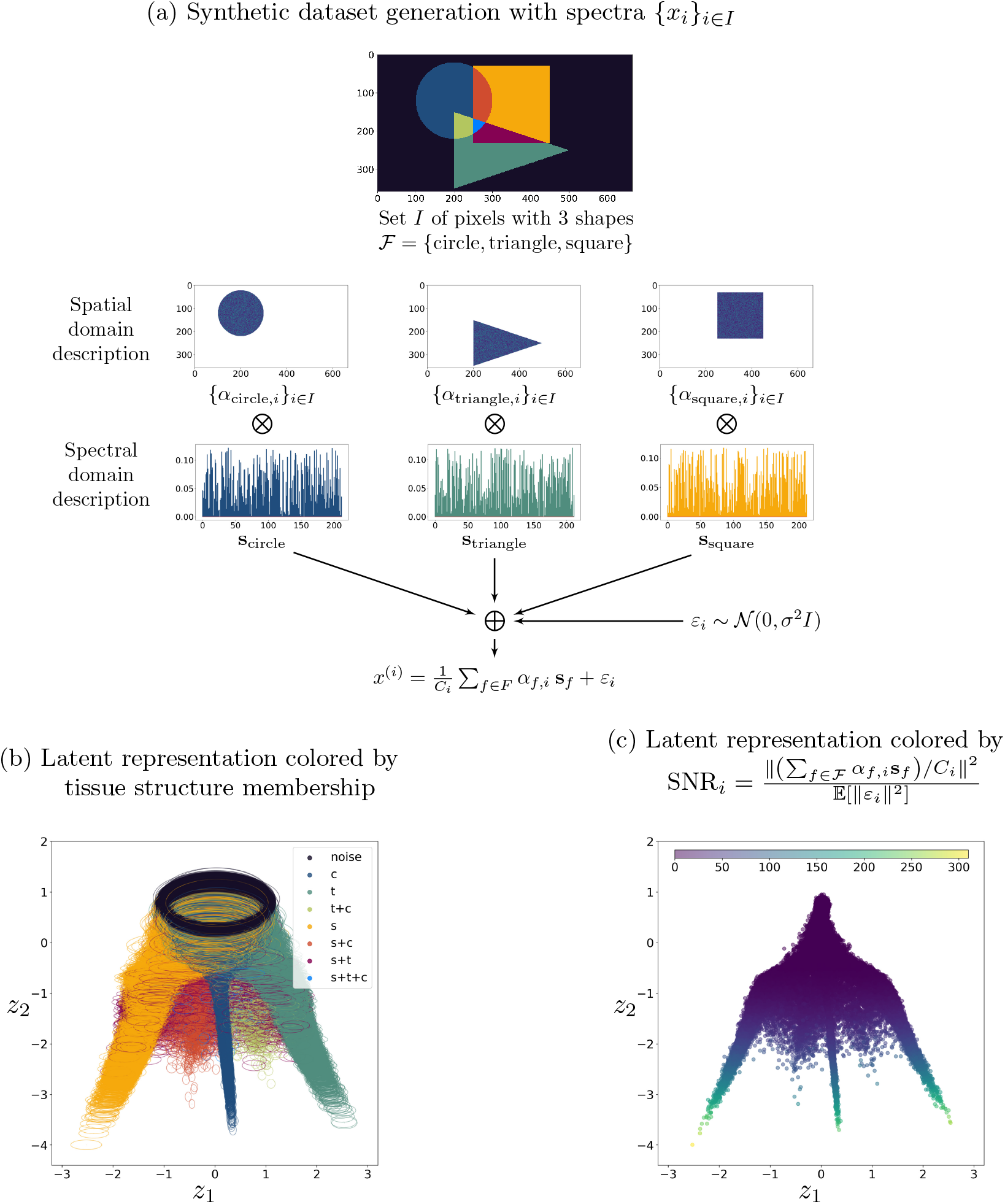
Workflow applied to case study 2’s synthetic dataset. (3a) Generated using three spatial ‘tissue structures’: ‘c’ (circle), ‘t’ (triangle), ‘s’ (square). Each shape has a distinct spectral signature, scaled and averaged to give individual pixel spectra. (3b) Latent representation with standard-deviation ellipse per spectrum, colored by its ‘tissue structure’ and ‘noise’ pixels not a member of any. Qualitatively different signatures (‘c’, ‘t’, ‘s’ legs) separate along the *z*_1_-axis. Pixels that combine shapes lie inbetween the corresponding legs, demonstrating *z*_1_’s interpretive power. (3c) Latent representation with mean location per spectrum, colored by its SNR_*i*_. SNR/signal strength separates along the *z*_2_-axis. Also, 3b‘s ellipse size, *i.e*., uncertainty, correlates with 3c‘s SNR, *i.e*., molecular abundance.

After applying our approach, a *β*-VAE-supplied latent space was obtained. Figure 3b shows the dataset’s latent representation in that space, using a standard-deviation ellipse per spectrum, emphasizing variance. Figure 3c shows the same representation, using the mean of each spectrum, emphasizing location. To examine where different content signatures are encoded in this space, each pixel’s ellipse in Figure 3b is colored according to the shape, or ‘tissue structure’, it belongs to (color-matched to Figure 3a). Qualitatively different signatures, *i.e*., shape contents, were separated along the *z*_1_-axis. Moreover, pixels associated with multiple shapes, *f*_1_ and *f*_2_, were lying in between the latent ‘legs’ reporting pure *f*_1_ and *f*_2_ pixels, hinting at the *z*_1_-axis’ advanced data interpretation potential. To explore where quantitative strength-reporting variation is encoded, Figure 3c colors each pixel according to its SNR_*i*_-value. This showed that SNR, representing molecular abundance, was encoded along the *z*_2_-axis (lower-signal-strength pixels at the top, high-signal-strength pixels at the bottom). The link between lower *z*_2_-values and increased SNR is also apparent in Figure 3b: uncertainty-reporting ellipses become smaller towards the bottom and pure-noise pixels with large variance localize to the top. This synthetic dataset showed that, without deliberate steering, the *β*-VAE-produced latent representation tends to disentangle two biologically relevant factors: signal content (reporting molecularly-distinct tissue structures) is modeled almost independently from signal strength (reporting molecular abundance). This case study confirms our workflow’s tendency to develop SiSAL spaces, at least for measurements that fit the signal model in section 4.3.

For comparison, traditional UMAP and t-SNE dimensionality reduction were applied to this dataset as well (Appendix F). The results varied strongly with hyperparameter choices. However, none of their latent axes cleanly separated signal content from signal strength, an important property offered by SiSAL spaces.

### 2.3 SiSAL spaces enable a novel algorithm to reveal distinguishing molecular features

In Figure 2, the *z*_1_-axis separated case study 1’s measurements by relative molecular content. Subsequently, the KDE-estimated latent density (Figure 2b) allowed latent ROIs to be defined (Figure 2c). When projected back onto tissue, we discovered that each latent ROI associated with a particular tissue structure (Figure 2d), presumably corresponding to different functional tissue units (FTUs) that constitute the renal nephron. This presumption can be independently verified: previous work [25] established a convolutional neural network that recognizes renal FTUs from autofluorescence microscopy images. Since these HuBMAP IMS-datasets include associated autofluorescence images, we obtained for each IMS pixel in case studies 1,3-5 an AF-based FTU label, reporting glomeruli (GL), proximal tubules (PT), distal tubules (DT), collecting ducts (CD), or thick ascending limb (TAL). If we color our IMS-based latent representation using these AF-based labels (Figure 4a), our presumption that different latent density ‘legs’ correspond to distinct FTUs is confirmed. For example, according to autofluorescence microscopy, the orange-colored ROI in Figure 2c consisted nearly exclusively of glomerulus-reporting pixels in Figure 4a.

**Fig. 4:**
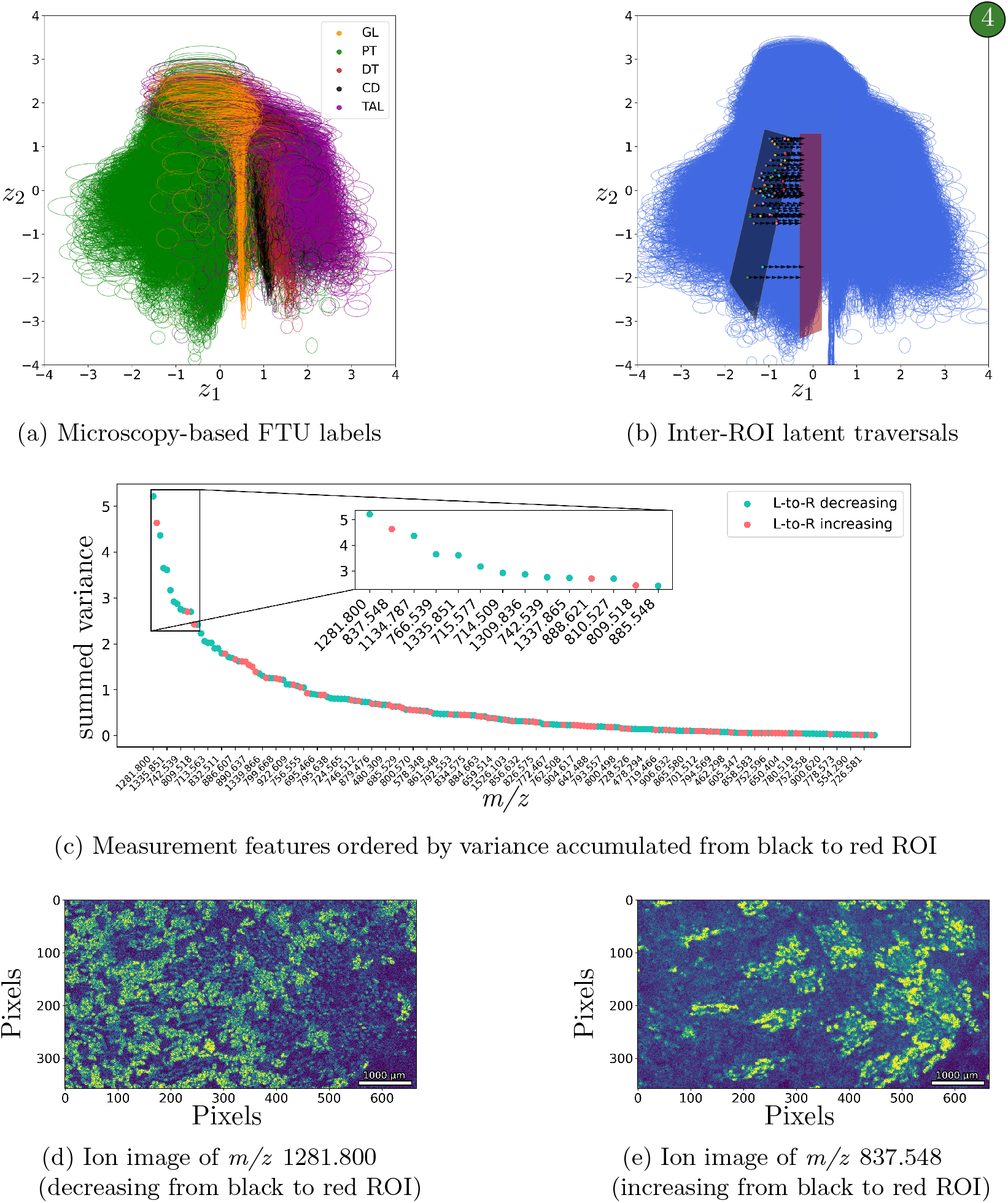
Workflow’s step 4 applied to case study 1’s human kidney IMS data. 4a Microscopy labels all green pixels as proximal tubules (PT), but our IMS-based latent space reveals heterogeneity, suggesting multiple substructures. Figure 2b also suggests at least two latent density ‘legs’ or molecularly distinct subsets within PT-labeled pixels. (4b) Guided by this, black and red ROIs were drawn to separate these substructures. (4c) Algorithm 1 ranks *m/z* -features by latent variance accumulated between ROIs, identifying *m/z* 1281.800 (a glycosphingolipid) and 837.548 (a PI) as the strongest differentiators, with opposite abundance trends. (4d-4e) Ion images confirm distinct localizations of these features with little overlap. This indicates a true chemistry-driven subdivision of PT, not captured by autofluorescence microscopy.

After independent confirmation that our latent representation captured biologically meaningful structure and establishing a link to the measurement space’s spatial domain, the need to connect latent findings also to the measurements’ spectral domain became compelling: it would facilitate interpretation in terms of molecular features. To address this challenge, we developed a novel comparative-latent-traversal algorithm (Algorithm 1) that uses the SiSAL structure to identify specific measurement features that differentiate latent ROIs. This process constitutes the final step of our workflow (Figure 1b), and is demonstrated in Figures 4b-4e. We illustrated the utility of Algorithm 1 by investigating heterogeneity found within the green-labeled pixels (Figure 4a). AF-based labeling considered all green pixels as reporting PT. However, based on IMS-measurements, our workflow suggested that Figure 4a‘s green latent area contains more than one ‘leg’, implying subgroups within the PT-related pixels. In fact, Figure 2b‘s KDE representation already revealed at least two latent density ‘legs’ or qualitatively different molecular signatures among the PT-labeled pixels. These findings demonstrate the workflow’s capability to detect known AF-labeled kidney structures and to further subclassify these structures based on their molecular content. This approach serves as a valuable hypothesis-driven detection tool for human kidney tissue exploration. To determine the molecular features responsible for differentiating these potential subdivisions of PT, we define the black and red ROIs and run Algorithm 1 between them. Figure 4b shows several of Algorithm 1‘s latent space traversals, oriented along the content-differentiating axis of the SiSAL space (here, *z*_1_), from left (black) to right (red). The traversals cover different locations along the SiSAL space’s

SNR-reporting axis (here, *z*_2_) to avoid accumulating variance specific to only certain signal strengths. Algorithm 1 ranks the measurement space’s features from most to least latent variance accumulated (Figure 4c). In this case, the two most varying molecular features were *m/z* 1281.800, a glycosphingolipid, and *m/z* 837.548, a phosphatidylinositol (PI). Since the algorithm captures features’ average intensity-change while traversing, it automatically determines for which ROI each molecular feature exhibits increased abundance. Specifically, *m/z* 1281.800 is increased in the black ROI and decreases towards the red ROI, while *m/z* 837.548 exhibits opposite behavior. This SiSAL-provided observation is confirmed when we retrieve the ion images for the two algorithm-suggested features from the 212 available (Figs. 4d-4e). The minimal spatial overlap of each uniquely localized lipid species dispels the idea that these lipids report the same FTU. This instead suggests a genuine, lipid profile-driven subclass of the PT that is distinct from classifications derived from AF microscopy.

### 2.4 Biological validation of SiSAL-based findings

Although confirmed in terms of chemical variation, determining whether the suggested PT-subdivision is genuinely biological in nature required further validation. To avoid dataset-or patient-specific observations, we applied our workflow to three additional IMS-measurements of human kidney tissues from different patients: HuBMAP VAN0063-RK-1-41, VAN0042-RK-1-31, and VAN0049-RK-1-35 (case studies 3-5, respectively). Case study 3’s results are shown in Figure 5, and case studies 4 and 5 are provided in supplementary Figures B1-B4. All three datasets yielded a SiSAL space (Figures 5a,5b) that clearly separated microscopy-labeled FTUs (Figure 5c), confirming that similar biology is captured across patients. As in case study 1, all three KDE representations also suggesed a PT-subdivision (Figure 5d), *i.e*., substructures within the green pixels. This consistency across patients, without prior guidance, is a testament to the robustness and reproducibility of our workflow.

**Fig. 5:**
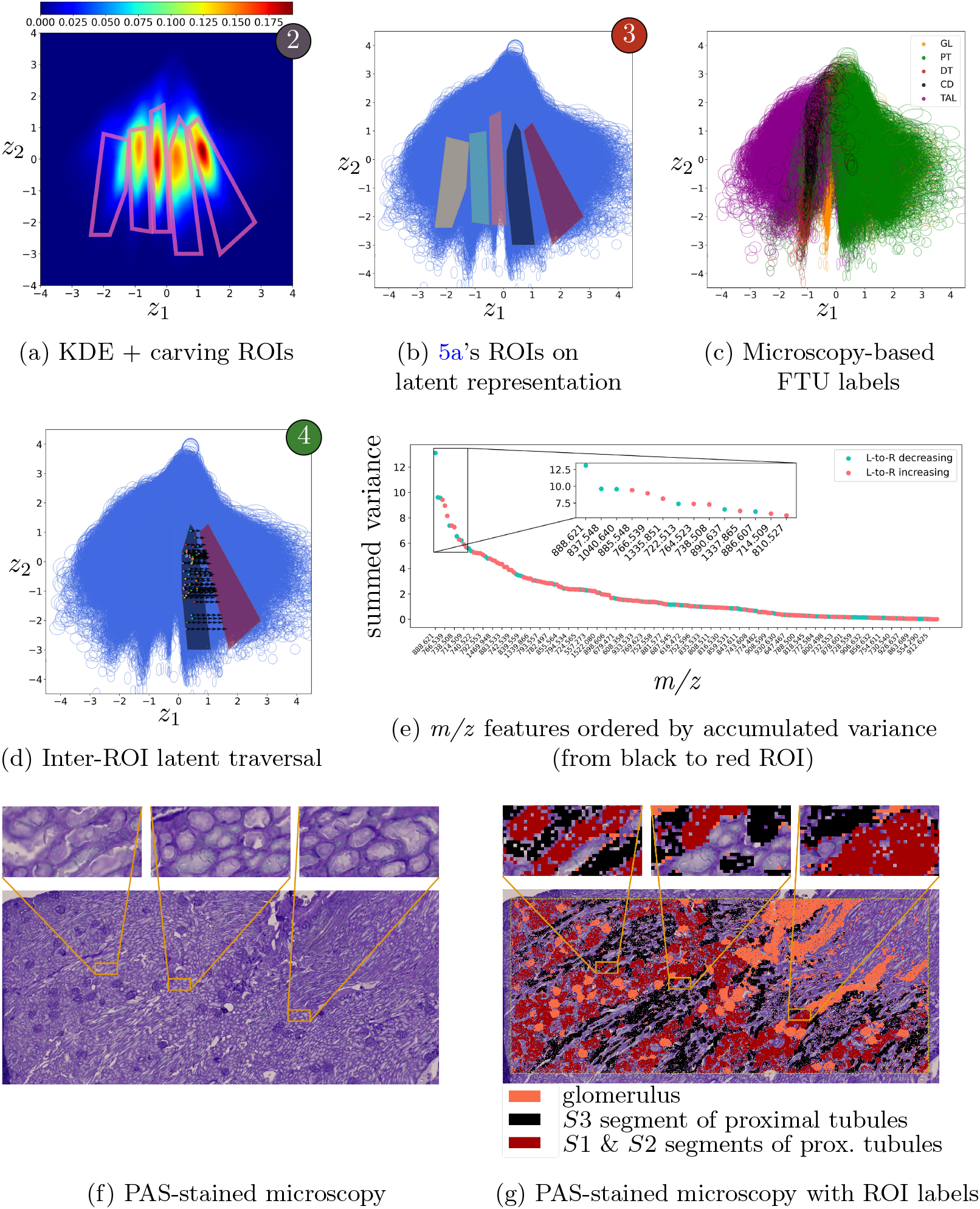
Workflow applied to case study 3’s human kidney data (VAN0063-RK-1-41). Figure 5a, 5b, and 5d report steps 2, 3 and 4. The low-dimensional representation of this dataset demonstrates a SiSAL structure: elongated abundance-reporting striations along one axis and the other axis separating these signatures as chemically distinct. Figure 5c colors the latent space using microscopy-based FTU labels, independently confirming that the found latent ‘legs’ correspond to different biological structures. The black and red ROIs are defined to examine subdivisions among green PT-reporting pixels. Our comparative-latent-traversal algorithm reveals (5e) which *m/z* -features vary most between these ROIs. In 5f and 5g, we map our findings to PAS-stained microscopy for biological interpretation.

First, we examined within the spatial domain whether the observed PT-subdivision aligned with known kidney morphology. We used Periodic acid–Schiff (PAS)-stained microscopy images from the same tissue sections as those used for IMS (Figure 5f). PAS is a modality used by renal pathologists to ascertain morphology and analyze lesions that can indicate specific diseases. Using these images, expert kidney pathologists found that the IMS-defined black ROI is associated with the S3 segment of the PT, while the red ROI correlates with the remaining S1 and S2 segments of the PT. Similar observations were seen in case studies 4-5 (Appendix B). Subsequently, our comparative-latent-traversal algorithm enabled us to explore the spectral domain as well. Algorithm 1 performed latent traversals between the black and red ROIs along the SiSAL space’s content-reporting axis (Figures 5d,B1d,B3d), summing up variance experienced by each measurement space feature while traversing. Hereby, Algorithm 1 effectively uncovers molecular differentiators between these latent ROIs. In descending order of traversal-accumulated variance, top measurement space features will tend to be those that change the most between the black and red ROIs, *i.e*., they report molecular features that are most different between the S3 and S1+S2 PT segments, respectively. Case studies 3 and 4 ran the algorithm from the black to red ROI, and case study 5 from red to black. Assuming biological consistency, in all case studies, the same molecular species should come to the forefront, with *m/z* -features that increase in case studies 3 and 4, decreasing in case study 5. Figures 5e,B1e,B3e indeed exhibited this behavior, verifying Algorithm 1‘s functioning.

Among the top-14 *m/z* -features (Figures 5e,B1e,B3e), we observed dataset-specific differentiators (*e.g*., *m/z* 801.677 and 1040.640), but also molecular features that differentiate S3 from S1+S2 across all three case studies (*e.g*., *m/z* 714.509 and 888.621). The former is not unexpected, particularly for human tissue studies, as each patient is distinct. The common findings, however, demonstrate that our workflow can pull nuanced, consistent observations from high-dimensional measurements, even in the presence of noise and inter-patient variation. Moreso, the molecular features suggested by our workflow appear consistent with known kidney biology. For example, polyunsaturated phosphatidylethanolamine (PE) species are known to regulate Na+/K+-ATPase activity [26]. Consistent with this, PE(34:2) (*m/z* 714.509 in Figure 4c,5g,B1,B3), PE(36:4) (*m/z* 738.508 in Figure 4c,5g,B1,B3), PE(38:5) (*m/z* 764.523 in Figure 5g,B3), and PE(38:4) (*m/z* 766.539 in Figure 5g,B3) were highly correlated with the S1+S2-region. Phosphatidylserine PS(38:4) (*m/z* 810.527 in Figure 4c,5g,B3) is even known to exclusively associate with the S1+S2-region. Besides confirming previous findings, our workflow also proposes new hypotheses. For example, we found phosphatidylinositol PI(38:4) (*m/z* 885.548 in Fig 4c,5g,B3) also differentiated the S3 from the S1+S2 segments of the PT, localizing to S1+S2 PTs. This colocalization of PI(38:4) with SGLT2 inhibitors in the S1/S2 segments suggests that PI(38:4) may also influence SGLT2 activity, potentially through the PI3K signaling pathway. More elaborate biological interpretation is provided in supplementary section E.

## 3 Discussion

Unsupervised dimensionality reduction has become key to analyzing complex, highdimensional molecular imaging data. Low-dimensional representations provided by, *e.g*., PCA or UMAP are commonly used for qualitative visualization or as a basis for clustering. However, interpreting these representations is oftentimes challenging. Their latent axes do not necessarily capture biology-relevant aspects, such as molecular composition or abundance. They also often lack an explicit connection back to the measurements, making it difficult to identify molecular changes that correspond to differences in latent location.

Our approach overcomes these challenges by learning a directly interpretable latent representation of high-dimensional molecular imaging data. Using a *β*-VAE, we not only project to a lower-dimensional representation, but also learn a generative model that explicitly captures uncertainty. This yields a latent space that separates signal strength from signal content, referred to as a signal strength aware latent (SiSAL) space. Although unsupervised, the consistently recurring SiSAL structure suggests that this approach captures fundamental properties of IMS data. This probabilistic exploration is further enhanced by our KDE-based segmentation of latent ROIs. In human kidney examples, SiSAL spaces not only revealed canonical features, such as glomeruli, but also previously-unrecognized molecular heterogeneity within proximal tubule segments. Translating such discovered latent ROIs into molecular features responsible for differentiating them is crucial for biological interpretation. Therefore, we introduced a novel comparative-latent-traversal algorithm that converts differences between latent regions into ranked molecular features. This provides a direct link between abstract locations in the latent space and specific molecular features, enabling biologists and clinicians to interpret low-dimensional data representations in terms of molecular species.

This work opens up new possibilities for analysis in spatial biology. Instead of being solely descriptive, *β*-VAE-derived SiSAL spaces provide structured, low-dimensional data representations that support interpretation, discovery, and hypothesis-generation in large-scale spatial omics studies.

## 4 Methods

### 4.1 IMS data and preprocessing

An IMS experiment records *n* pixels, each reporting a full-profile mass spectrum. After preprocessing and peak-picking, each of these *n* mass spectra is reduced to a vector of *m* scalar ion intensity values, reporting the abundances of *m* distinct ion species at that spectrum’s location in the tissue. We denote an IMS dataset 𝒟 as a set of *n* spectra, *i.e*., 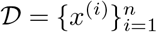, each reporting *m* features, *i.e*., *x*^(*i*)^ ∈ ℝ ^*m×*1^. As our workflow requires training a model, the dataset 𝒟 is split into a training set 𝒟^*′*^ and a validation set 𝒟 \ 𝒟^*′*^ through random sampling, approaching a 80*/*20 training/validation split among the available spectra. This separation helps prevent overfitting during training. We monitor performance on the validation set, halting training after no improvement for three consecutive epochs. The data is standardized, *i.e*., mean-subtracted and scaled per-feature to a standard deviation of 1, such that

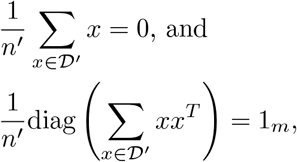

where *n*^*′*^ = |𝒟^*′*^| is the number of spectra/pixels in the training set, diag(*A*) is the vector formed by the diagonal elements of matrix *A*, and 1_*m*_ ∈ ℝ^*m×*1^ is a vector consisting only of 1s. The standardization is based on the training data 𝒟^*′*^ to avoid leakage during model validation. Standardizing the data ensures that each feature is centered around 0 and has a variance of 1, preventing ion species’ different absolute intensity ranges from biasing subsequent analysis and ensuring that every feature, and thus ion species, has a chance to be equally impactful.

### 4.2 *β*-VAE-based workflow

The workflow we developed enables a probability-aware (rather than deterministic) unsupervised exploration of an IMS dataset, learning generative factors that underlie the measured spectra. In the context of IMS-based characterization of the molecular content of a tissue section, finding generative factors for a dataset amounts to distinguishing different sources of variation within the measurements of the IMS experiment. Some of these sources of variation can be biological in nature, while others can report technical variation introduced at the sample preparation or instrumental level. Once found, these generative factors describe a lower-dimensional latent space in which the IMS data can be represented compactly. Every empirically acquired mass spectrum can be represented as a combination of generative factors and thus corresponds to a specific location within the latent space. Since the latent space is variational in nature, it allows for a probability density-based interpretation of its structure. This facilitates the discovery and delineation of high-density latent regions-of-interest (ROIs), which tend to correspond to specific biological structures in the measured tissue. As such, the generative latent representation of an IMS dataset will typically tend to report and confirm known tissue structure. More importantly, it can also suggest new, hereto unrecognized structures within the tissue, purely on the basis of the molecular content reported by IMS. Finally, through a novel comparative-latent-traversal algorithm, our workflow goes beyond merely suggesting potential tissue structures and instead characterizes them in terms of the molecular features, *i.e*., ion species, responsible for their differentiation from other structures. The algorithm permits for the differences between ROIs in the latent space to be cast back onto the original measurement space features, such that the ion species differentiating between generative factor-delineated tissue structures can be interpreted in both a chemical and a biological context.

The workflow consists of four steps:

- **(Step 1)** A *β*-VAE is trained to find a low-dimensional latent space to represent the provided data, with the dimensionality of the latent space specified by the user. The variational nature of this latent space, together with the pressure exerted by the *β* parameter in the objective function, ensures that the axes describing the space implicitly encode generative factors for the approximated dataset. Interestingly, we can observe that the combination of the variational requirements to construct this space and the nature of mass spectral measurements tends to yield generative factors that separate out absolute signal strength or SNR from relative signal patterns and their qualitative differentiation. For brevity, we will refer to such a space as a SiSAL space going forward. In this paper, we specify latent spaces to have a dimensionality of two. While any positive integer number of latent dimensions is valid and can be pursued with the workflow described here, in our case studies we restrict ourselves to two latent dimensions because it allows for a clear demonstration of SiSAL spaces’ tendency to separate along a qualitative axis and a quantitative axis, while still being straightforward to interpret by humans in a plot. At least for mass spectral data, step 1 is demonstrated to be able to yield SiSAL spaces, and we will show their separating characteristics to be extremely valuable for data interpretation.
- **(Step 2)** A latent density function is estimated from the *β*-VAE-supplied latent representation, using KDE with a Gaussian kernel. The density representation reports a rasterized function across the latent space and reveals nuances that are challenging to observe directly in the level-set and point-cloud-based representations supplied by step 1.
- **(Step 3)** As the density-based view into the latent space brings out high-density areas clearly, delineation of distinctive latent ROIs as polygons becomes possible. Since the latent density representation is used here as a discovery tool to discern both known and previously unknown structures within the tissue, the ROI delineation is performed manually and is human-driven. However, the latent density space offers opportunities for automated latent ROI determination as well, particularly if a SiSAL-like structure is also present. Each of the ROIs defines a latent space subarea corresponding to the probabilistic structure underlying the dataset. However, by means of the mass spectral measurements (or pixels) that fall within its polygon when mapped to the latent space, every ROI also implicitly connects to a set of pixels and, thus, a spatial subarea within the measured tissue. This connection between the latent space and the spatial domain of the measurement space offers an advantageous duality. It is easier to discover and delineate ROIs in the latent space than in the tissue’s measurement space. The latter, however, reveals which tissue structures a high-density ROI corresponds to, and in many cases, this will separate out genuinely biological structures such as morphological tissue areas, functional tissue units, cell types, and cell states. Furthermore, if the latent space is SiSAL-like, this can be taken even further: it allows one to make tissue structure delineations independent of signal strength.
- **(Step 4)** While step 3 delivers implicit molecular differentiation between different latent ROIs and their corresponding tissue structures, it does not reveal the measurement features responsible for that differentiation. Since that information is locked into the combination of the *β*-VAE encoder and the established latent space, we need a means of translating latent space differentiation back into differentiation in the original measurement space. To accomplish this, we develop a novel comparativelatent-traversal algorithm that utilizes the *β*-VAE decoder and the qualitative axis of the SiSAL space to learn the measurement space features, *i.e*., ion species, most responsible for the differentiation between two latent ROIs.

Subsequent sections provide more details for each step in the workflow.

#### 4.2.1 Step 1 – Learning a variational latent space by *β*-VAE

While common manifold learning techniques, such as t-SNE and UMAP, can deliver latent spaces that represent a given dataset sufficiently well in terms of approximation, this does not necessarily mean the provided latent space encodes relevant structure. Furthermore, such latent spaces typically do not provide a concept of uncertainty, making it difficult to assess how measurement-space variance corresponds to latent-space variance. In step 1, we seek to address both issues by learning a probabilistic rather than a deterministic latent space. Instead of seeking latent axes that are merely dimensionality-reducing, the latent space we strive for is optimized to yield axes that can serve as generative factors for the measured data, and is shaped to inherently accommodate uncertainty around the measurements. Specifically, we are interested in finding an estimate of the underlying probability distribution *p*(*x*) that generates the samples in 𝒟. Moreover, we aim to utilize this distribution to explore and draw meaningful conclusions about the data itself.

The most common approach to obtain *p*(*x*) is to use a family of distributions 𝒫 = {*f*_*θ*_ | *θ* ∈ Θ }, where *θ* is a particular set of parameter values from the family’s parameter space Θ, and to pick the parameter value set that fits the data best. However, for distribution families with a parameter space that is rather large, it becomes harder to draw meaningful conclusions about the data directly from those parameters. On the one hand, when the parameter space grows, it becomes harder to understand what a particular parameter changes in the overall distribution. On the other hand, with a small number of parameters like in a (unidimensional) Gaussian family, for example, we can only capture one mode of the data, which is insufficient for most multidimensional measurement types.

VAEs [16] circumvent this issue by assuming that the learned distribution can be as complex as necessary, but that it depends only on a few factors represented by a low-dimensional vector *z* ∈ ℝ^*d*^ with *d* ≪ *m*. This induces a distribution *p*_*θ*_(*x*| *z*) where *θ* is the vector of all parameters necessary to parameterize the distribution. With this model, a sample *x*^(*i*)^ from the distribution is obtained by:

1. sampling *z*^(*i*)^ from the prior distribution *p*(*z*); and
2. sampling *x*^(*i*)^ from the conditional distribution *p*_*θ*_ *x*|*z*^(*i*)^ .

In our study, we choose a standard normal prior *p*(*z*) = 𝒩 (0, *I*), where *I* is the identity matrix, and a Gaussian likelihood *p*_*θ*_(*x* | *z*) = 𝒩 (*µ*_*θ*_(*z*), *I*). The mapping *µ*_*θ*_(*z*) : ℝ^*d*^ → ℝ^*m*^ denotes the mean of the likelihood function at *z* and is parameterized by *θ*. Note that for complex *µ*_*θ*_(*z*), such as ones provided by neural networks, the density of *x*

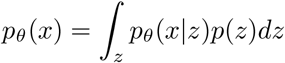

is not necessarily Gaussian and is generally intractable. This means that we cannot find an analytical solution to this integral [16]. One could use Bayes’ rule to find the posterior distribution:

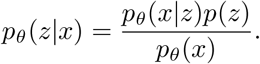

However, with the normalizing constant *p*_*θ*_(*x*) being generally intractable, *p*_*θ*_(*z* | *x*) is also generally intractable. The VAE solves this by additionally learning an approximate posterior distribution *q*_*ϕ*_(*z*|*x*) ≈ *p*_*θ*_(*z*|*x*), parameterized by *ϕ*. In this work, we choose to model *q*_*ϕ*_(*z*|*x*) using the family of Gaussian distributions with a diagonal covariance matrix. Thus, *q*_*ϕ*_(*z* | *x*) = 𝒩 (*µ*_*ϕ*_(*x*), Σ_*ϕ*_(*x*) = diag(*σ*_*ϕ*_(*x*))) where diag(*a*) denotes the diagonal matrix holding the entries of the vector *a* on its diagonal. Forcing a diagonal covariance matrix ensures that the learned factors *z* are independent. The subscript *ϕ* denotes the set of all parameters needed to parameterize the mean *µ*_*ϕ*_(*x*) and variance *σ*_*ϕ*_(*x*) mappings. More concretely, in the context of neural networks, *θ* and *ϕ* refer to the complete set of weights that configure the network.

By learning the functions *p*_*θ*_(*x*| *z*) and *q*_*ϕ*_(*z*| *x*), we create probabilistic mappings between the native space of the data (ℝ^*m*^) (the measurement space) and the space of the generative factors (ℝ^*d*^) (the latent space). As *q*_*ϕ*_(*z* |*x*) encodes the data into a probabilistic latent space, it is commonly referred to as a probabilistic encoder. Reciprocally, *p*_*θ*_(*x*| *z*) is a mapping from the latent space to the native measurement space, and it is called a probabilistic decoder. As mentioned above, in this paper we will focus on the case *d* = 2. A visual representation of this is shown in Figure 1a.

The VAE objective loss function corresponds to the negative evidence lower bound (ELBO) [27]. Over all *n* samples, this is:

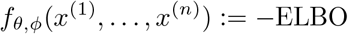

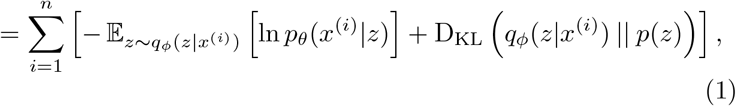

where, for two densities *p*_1_ and *p*_2_, 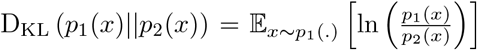 is the Kullback-Leibler divergence between the two densities (see Theorems 2 and 3).

##### What is the ELBO?

As its name suggests, ELBO is a lower bound on the evidence:

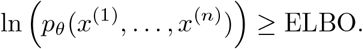

From (1), we see that *f*_*θ,ϕ*_ can be separated in two objective functions:

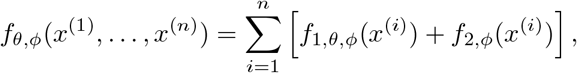

with 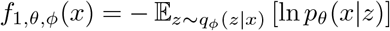 and *f*_2,*ϕ*_(*x*) = D_KL_ (*q*_*ϕ*_(*z*|*x*) || *p*(*z*)).

Minimizing *f*_1,*θ,ϕ*_(.) forces the distribution *q*_*ϕ*_(*z*|*x*) to concentrate on the *z*’s that maximize *p*_*θ*_(*x*|*z*), *i.e*., it forces *q*_*ϕ*_(*z*|*x*) to move to *z*’s that yield a large likelihood *p*_*θ*_(*x*|*z*). This part of the objective function can be seen as the dataset-approximating part, since the *z*’s with a large likelihood *p*(*x*|*z*) can be used to retrieve *x*.

Minimizing *f*_2,*ϕ*_(.) forces the distribution *q*_*ϕ*_(*z*| *x*) to remain close to the standard normal prior *p*(*z*). This part of the objective function is responsible for the structure in the latent *z*-space (and thus also its interpretability). In Theorem 3 of Supplementary section D, we recall a standard argument for this term forcing the mean and covariance matrix of *q*_*ϕ*_(*z*|*x*) to be close to the zero vector and the identity matrix, respectively. For *x*^(*i*)^ ∈ *D*, this forces a common structure on the posterior distributions *q*_*ϕ*_(*z*|*x*^(*i*)^).

##### Importance of β in the β-VAE

The *β*-VAE [17] is a modification of the VAE that adds a hyperparameter *β* ∈ ℝ^+^ to the objective function:

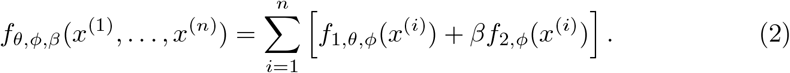

With *β >* 1, we force additional importance onto the term *f*_2,*ϕ*_(.) of the objective function in (2). This term acts as a regularization term, preventing the latent representation *z* ∼ *q*_*ϕ*_(*z* | *x*) from wandering too far from the origin and exhibiting too small a variance (*e.g*., uninformative factors *z*). We observe in the Results section that when this method is combined with mass spectra of an IMS experiment, it tends to recurringly yield a common structure of the latent space, which we refer to as a SiSAL space.

Given that interpretability is a central objective in our work, the extraction of informative latent axes is essential. Since *β*-VAE’s *β* provides a means of enforcing such axes, we use the *β*-VAE rather than the standard VAE to establish the probabilistic latent space for step 1 of our workflow. Practically, step 1 uses the training and validation datasets specified above to train two neural networks: the probabilistic encoder, implementing *q*_*ϕ*_(*z*|*x*) and shown on the left in Fig. 1a, and the probabilistic decoder, implementing *p*_*θ*_(*x*| *z*) and shown on the right in Fig. 1a. The specific architecture of the neural networks is provided in Supplementary section A. During the training process, a specific *β*-value will be used to drive the optimization of the weights. In terms of setting the value of *β*, its particular value is not pertinent as long as sufficient pressure is exerted such that informative factors *z* appear. Specifically, in all case studies in this work, we manually increased *β* until an informative latent space became available, and consistently this space tended towards the qualitativeversus-quantitative separation characteristic of a SiSAL space. Since subsequent steps in our workflow carve out ROIs of high latent density, any latent space that separates high-density areas sufficiently well to perform that carving is sufficient for the remainder of our workflow. As such, there is not one particular *β*-value to consider as long as we use a value that yields an informative latent space. Nevertheless, the *β* in *β*-VAE is essential as the standard VAE does not necessarily deliver the latent structure we need for the subsequent steps in our exploratory workflow.

Once *ϕ* and *θ* have been optimized, *i.e*., respectively the encoder and decoder are trained, we can plot the level sets of *q*_*ϕ*_(*z*|*x*) for all *x* ∈ 𝒟. Since we chose a Gaussian distribution for *q*_*ϕ*_, its level sets correspond to ellipses. In most visualizations of the obtained latent space (Figs. 1b, 2a, 3b, 4a, 2b, 5b-5d, B1b-B1d, and B3b-B3d), we use the ellipse associated with one standard deviation around the mean as a representation tool:

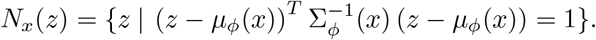

This ellipse allows us to visualize the mean and the standard deviation on the *z*-axes for each sample *x*^(*i*)^’s latent distribution 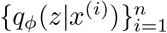. The ellipse not only models sample *x*^(*i*)^’s mean location in the *z*-space, *µ*_*ϕ*_(*x*), but it also reports the uncertainty around that mean location. This notion of uncertainty answers a key requirement and is a strong differentiator from other dimensionality reduction approaches, and we use it specifically in our workflow’s step 4, where Algorithm 1 accumulates this latent uncertainty when performing a latent traversal between ROIs. An example of the latent space representation with *d* = 2 of an IMS dataset, where an ellipse depicts the uncertainty around a pixel’s mapping into latent space, is shown in Figure 2a. While some structure is visible, the depicted cloud of ellipses makes interpretation and delineation of latent ROIs difficult, and step 2 is needed to make this informative latent space human-perusable.

#### 4.2.2 Step 2 – Estimating latent density by KDE

After establishing an informative *z*-space in step 1, we want to draw meaningful insight from the samples’ latent distributions }*q*_*ϕ*_(*z*| *x*^(1)^), …, *q*_*ϕ*_(*z* |*x*^(*n*)^)} in that space. For example, step 1’s *β*-VAE-driven latent space forces independence of its axes, *i.e*., factors. If measured data points *x*^(1)^, …, *x*^(*n*)^ share factors, this sharing should be reflected in their mean locations in the latent space. As such, the organization of samples’ latent distributions can reveal quite nuanced relationships between those samples in the native measurement space. However, for datasets with a large *n*, it is nontrivial to interpret the structure of that dataset’s latent representation from its level-set visualization (one standard deviation ellipse per pixel; example in Fig. 2a) or its point-cloud visualization (one mean location per pixel; example in Fig. 3c). We need an alternative view into the latent representation that is more effective at conveying the large amount of information present, that allows for direct human interpretation, and that facilitates straightforward manual or automated definition of latent ROIs in step 3. Step 2 pursues such an alternate view of the dataset’s latent representation by estimating the density throughout the latent space. Specifically, we examine samples *x*^(1)^, …, *x*^(*n*)^’s predicted means *U* = {*µ*_*ϕ*_ (*x*^(1)^), …, *µ*_*ϕ*_ (*x*^(*n*)^)} with *µ*_*ϕ*_ *x*^(*i*)^ ∈ ℝ^*d*^ (in our case studies, *µ*_*ϕ*_ *x*^(*i*)^ ∈ ℝ^2^) and their distribution.

Since *x*^(*i*)^ ∼ *p*(*x*) are samples from a distribution, the elements of *U* can also be viewed as samples from a distribution *h*(*z*). We want to understand the properties of that distribution. Applying a change of variable (see Theorem 1 of Appendix C), we find that

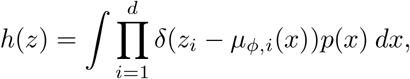

where *δ* is the Dirac delta function. Even if we knew *p*(*x*), this integral would still be intractable as the mapping *µ*_*ϕ*_(*x*) is performed by a neural network. Therefore, we need an alternative means of estimating the density of the *z*-space.

One solution is to use kernel density estimation (KDE), a parametric method, to find an estimate *ĥ*(*z*) of the distribution *h*(*z*). For a kernel function *K*, the kernel density estimate is

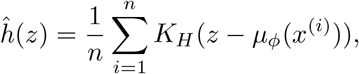

where *H* is a symmetric *d × d* matrix, referred to as the bandwidth of the estimate, and *K* is the kernel function. In this work, we choose *K* to be the multivariate Gaussian kernel *K*_*H*_ (*x*) = |det(*H*) |^−1*/*2^*K H*^−1*/*2^*x* . For the bandwidth, Scott’s rule of thumb [28] suggests picking a diagonal matrix *H* such that

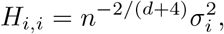

where 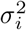 is the variance of the *i*-th feature. For all KDE-based latent density estimations throughout this paper, we use Scott’s rule of thumb.

Once estimated, we can use *ĥ*(*z*) to generate a rasterized representation of density across the latent space. This effectively functions as the alternative view we need into the latent representation (Figs. 2b, 5a, B1a, B3a). Contrary to the level-set and point-cloud representations, the density-based visualization reveals which regions in the latent space carry the most mass and it allows for straightforward delineation of high-density areas. In the Results section, we demonstrate that these high-density areas, which in the case of SiSAL spaces and IMS data recurringly take the form of high-density ‘legs’ oriented along the SNR-axis, tend to correspond to genuine biological structures within the tissue. This emphasizes the direct interpretability offered by step 2’s density-based latent representation of the data.

#### 4.2.3 Step 3 – Latent ROI definition & segmentation

Step 3 defines latent ROIs on the basis of the density-based dataset representation provided by step 2. As mentioned, the connection between features of the measurement space (*i.e*., the mass spectral domain) and the latent space is generally intractable: step 4 in our workflow addresses this challenge. However, the connection between locations in the measurement space (*i.e*., the spatial domain) and the latent space is more readily accessible. Pixels *x*^(1)^, …, *x*^(*n*)^ map to mean points *µ*_*ϕ*_(*x*^(*i*)^), …, *µ*_*ϕ*_(*x*^(*n*)^) in the latent space. If a latent ROI has been defined and a pixel *x*^(*i*)^’s latent mean *µ*_*ϕ*_(*x*^(*i*)^) falls within that ROI, we can consider that pixel a ‘member’ of that ROI. Taken together, the pixels that are members of an ROI implicitly define a spatial segment of locations within the measurement space that effectively functions as the spatial footprint of this ROI. It is this spatial mapping that we use to obtain Figures 2d, 5g, B1g, B2, B3g, and B4.

Defining a latent ROI can be a manual process, where a human investigator draws a region-of-interest within the *z*-space, using the dataset’s latent representation as a guide. ROI delineation can also be performed as an automated process, where an algorithm uses the dataset’s latent representation to define areas of interest in an automated way. Furthermore, the ROI definition process can be driven by prior knowledge (*e.g*., which latent structure corresponds to a certain FTU-of-interest?) or it can serve open-ended exploration (*e.g*., which tissue structures can be chemically discerned within this given IMS dataset?). As such, step 3 of our workflow can be implemented in different ways to serve either more targeted or more exploratory applications.

In this study, since we focus on open-ended exploration and we want to highlight potentially unexpected findings that this workflow can help uncover, we implement step 3 as a manual ROI definition. Specifically, the investigator ‘carves out’ ROIs by drawing a polygon in *z*-space, guided by step 2’s KDE-based estimate of high-density areas. Examples of such manually defined latent ROIs can be found in Figure 2b. Since the *z*-space is the same across the different visualizations, the ROI-polygons can be copied to other representations as needed. An example of such a transfer to the level-set representation of the same dataset is shown in Figure 2c. Using the spatial mapping described earlier in this section, we can now find out the spatial footprint of each ROI in the original measurement space. This, in turn, allows us to ascertain which biological tissue structure corresponds to which ROI. An example of this mapping is shown in Figure 2d. Besides manual ROI definition, there are also clear opportunities for algorithmic delineation of latent ROIs. In this paper, we focus on manual definition of ROIs because it allows us to highlight the human interpretability requirement we put forward at the start of developing this workflow. It also allows us to demonstrate how the workflow can be used in a human-in-the-loop manner, as a hypothesis-generating discovery tool whose findings can feed back into the wet-lab for follow-up experiments.

#### 4.2.4 Step 4 – Comparative latent traversal & identification of measurement-space features as differentiators

Step 4 takes the latent ROIs defined in step 3 and seeks to identify which measurement-space features differentiate the most between them. In the context of IMS-based characterization of organic tissue, step 4’s task amounts to taking biological structures captured by latent ROIs in step 3, and identifying which molecular species (*m/z* - features) are differentiators between those biological structures. Sometimes step 4’s output will confirm prior biochemical knowledge on the tissue at hand, sometimes step 4 will reveal a hereto unknown molecular differentiator. An example of the latter is shown in the Results section, revealing lipid species that differentiate S3 from S1+S2 PT segments in the human kidney.

As mentioned in the previous section, the link between the latent space and features in the measurement space (*i.e*., the mass spectral domain) is generally intractable. Therefore, we employ an alternative means of inferring differentiating features in the measurement space, namely through latent traversals. For this, we developed a modified latent traversal algorithm.

Latent traversals involve moving in unitary directions within the latent space and assessing concurrent changes in reconstruction. Since the factors in *z* are (nearly) independent, we can move in unitary directions by changing only one factor at a time. Latent traversals were already motivated in [17]. Here, we modify them to only traverse between two latent ROIs, making step 4’s results focus only on differentiators between the two corresponding tissue structures. Furthermore, we leverage the inherent separation of strength-versus-content offered by the SiSAL space to remove the influence of signal strength or SNR from the analysis. This is accomplished by letting traversals take place along the content-separating axis of the SiSAL space (specified to the algorithm by a unitary vector *e*), effectively amplifying content-related changes in step 4’s results. It also entails performing *n*_tr_ distinct traversals (in our case, *n*_tr_ = 50), each starting from a different random location along the signal strength-encoding axis of the SiSAL space, to attenuate SNR-related changes in step 4’s results. Overall, our comparative-latent-traversal algorithm computes the signed variance for each measurement space feature as we traverse from ROI to ROI.

Algorithm 1 describes our modified procedure. It requires two latent regions-of-interest *r*_1_ and *r*_2_. We take small steps in unitary directions to move from *r*_1_ to *r*_2_ (here, along the content-separating axis). At each step, we use our location in *z* and the decoder network, *p*_*θ*_(*x* | *z*), to sample a potential observation *x* in the native measurement space. We then compute the variance of these reconstructed observations, *i.e*., mass spectra, to identify the *m/z* -features that change the most. Examples of this algorithm’s output are shown in Figures 4c, 5e, B1e, and B3e.

##### Algorithm 1

Comparative latent traversal

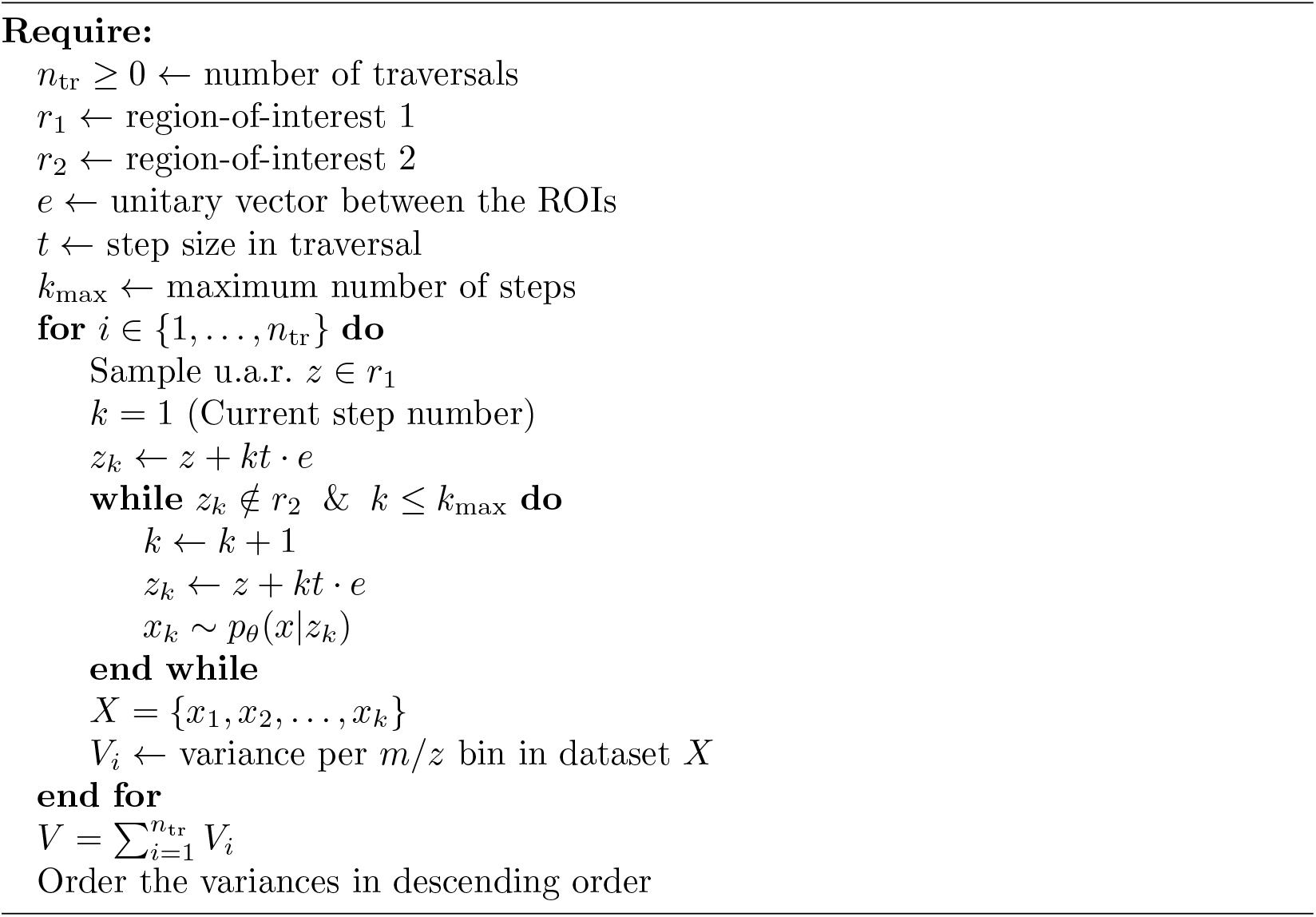

### 4.3 Synthetic dataset

For case study 2’s synthetic dataset, we design an image that contains a set of three overlapping shapes ℱ = {circle, triangle, square} (Figure 3a, top row), with *I* the set of pixels in that image. The shapes represent different tissue structures reporting different molecular content. For each shape *f* ∈ ℱ, *P*_*f*_ ⊂ *I* is the set of pixels that are part of shape *f* . Since the shapes overlap, ∩_*f*∈ℱ_ *P*_*f*_ ∅. For every shape *f*, a coefficient value is assigned to pixel *i*, representing that pixel’s signal strength or SNR:

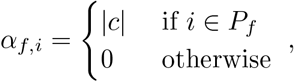

where 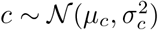. These spatial domain descriptions are shown in Figure 3a, middle row. Every shape *f* also has a unique deterministic spectral signature 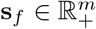 of norm one (Figure 3a, bottom row), with *m* the number of features per spectrum. Given the spatial and spectral signatures for each shape, a spectrum *x*^(*i*)^ is mixed for each pixel *i* ∈ *I*:

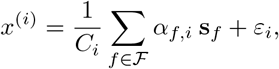

where *C*_*i*_ = ∑ _*f*∈ℱ_|*P*_*f*_ ∩ *i*| is the number of shapes that pixel *i* is part of, and 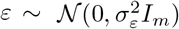 is a Gaussian noise perturbation. The distribution parameters for *c* and *ε* are provided in Appendix 2.2. If pixel *i* is only part of a single shape *f*, its mass spectrum *x*^(*i*)^ = *α*_*f,i*_ **s**_*f*_ + *ε*_*i*_ and its SNR measure is

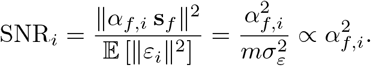

Thus, for the single-**s**_*f*_ case, pixel *i*’s SNR is proportional to the abundance coefficient 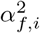.

### 4.4 Case studies

## 5 Acknowledgements

Research reported in this publication was supported by the National Institutes of Health (NIH)’s Common Fund, National Institute Of Diabetes And Digestive And Kidney Diseases (NIDDK), and the Office Of The Director (OD) under Award Numbers U54DK120058, U54DK134302, and U01DK133766 (J.M.S. and R.V.), by NIH’s Common Fund, National Eye Institute, and the Office Of The Director (OD) under Award Number U54EY032442 (J.M.S. and R.V.), by NIH’s National Institute Of Allergy And Infectious Diseases (NIAID) under Award Numbers R01AI138581 and R01AI145992 (J.M.S. and R.V.), by NIH’s National Institute On Aging (NIA) under Award Number R01AG078803 (J.M.S. and R.V.), by NIH’s National Cancer Institute (NCI) under Award Number U01CA294527 (J.M.S. and R.V.), and by the National Science Foundation Major Research Instrument Program CBET – 1828299 (J.M.S.). The research was furthermore made possible in part by grant numbers 2021-240339 and 2022-309518 (L.G.M. and R.V.) from the Chan Zuckerberg Initiative DAF, an advised fund of Silicon Valley Community Foundation. The results are in whole or part based upon data generated by the HuBMAP Program: https://hubmapconsortium.org. The content is solely the responsibility of the authors and does not necessarily represent the official views of the National Institutes of Health.

## Supplementary Information

### Nomenclature

*β-VAE*: *β*-variational autoencoder
*m/z*: mass-to-charge-ratio
*AF*: autofluorescence
*CD*: collecting ducts
*DT*: distal tubules
*ELBO*: evidence lower bound
*FTU*: functional tissue unit
*GL*: glomeruli
*HIS*: hyperspectral imaging
*HuBMAP*: Human Bio-Molecular Atlas Program
*IMS*: imaging mass spectrometry
*KDE*: kernel density estimation
*NMF*: non-negative matrix factorization
*PAS*: Periodic acid–Schiff
*PCA*: principal component analysis
*PT*: proximal tubules
*ROI*: region-of-interest
*SiSAL*: signal strength aware latent
*SMM*: spiked mixture model
*SNR*: signal-to-noise-ratio
*t-SNE*: t-distributed stochastic neighbor embedding
*TAL*: thick ascending limb
*UMAP*: uniform manifold approximation and projection
*VAE*: variational autoencoder

## Supplementary A Architecture and hyperparameters

### A.1 Model Architecture

Table A1 shows the VAE architecture for the different datasets. For all the datasets, we stopped training after no improvement on the test set for three epochs.

**Table A1:**
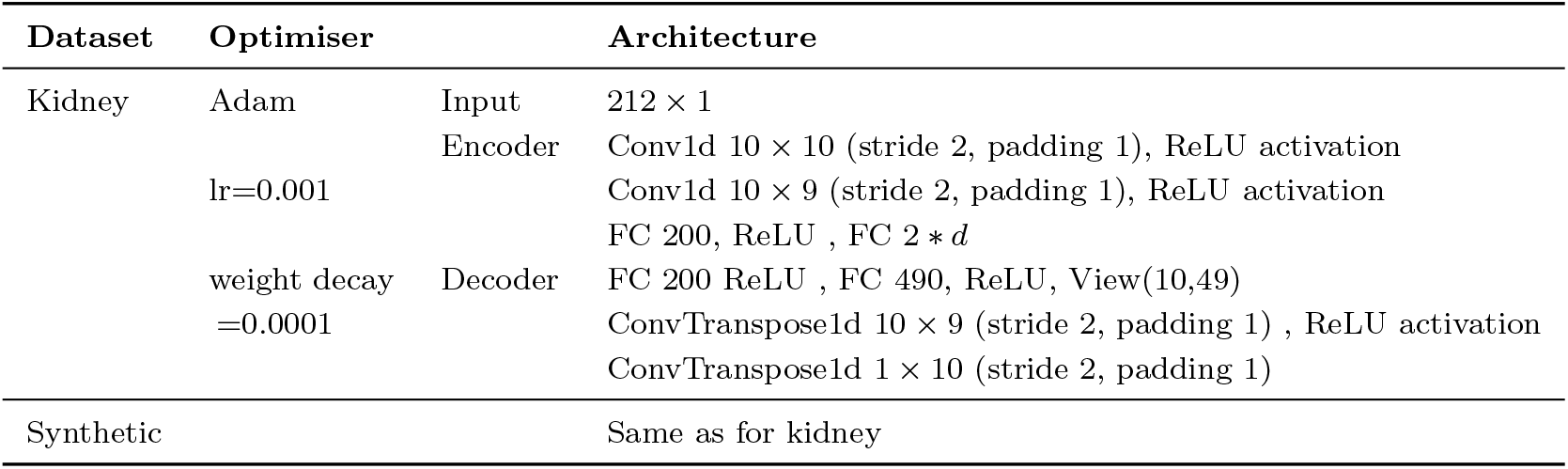
Details of model architectures. Conv1d(out channels, kernel size). Use of batch normalization

### A.2 Hyperparameter choices for section 2.2

The distribution parameters of 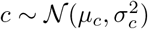 and 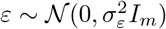 were:

- *µ*_*c*_ = 500
- *σ*_*c*_ = 500
- *σ*_*ε*_ = 10

## Supplementary B Additional datasets

We repeated the experiment of section 2.4 on two additional datasets from the HuBMAP project [22] (VAN0042-RK-1-31 and VAN0049-RK-1-35 in negative mode, see Table 1). Figures B1 (VAN0042-RK-1-31) and B3 (VAN0049-RK-1-35) highlight the same subdivision between proximal tubules. The accompanying PAS microscopy images also suggest in these additional cases that the black color corresponds to the S3 PT segment, while the red color corresponds to S1+S2 PT segment. For both datasets, full sets of the latent ROIs’ spatial segments are shown in Figures B2 and B4, respectively.

Note that, in Figure B1b, our workflow additionally suggests the presence of a yellow ‘leg’ that is not oriented along the SiSAL space’s SNR-reporting axis. Instead, its pixels seem locked at a fixed amount of (low) signal strength. When we cast the yellow ROI to the spatial domain using the method described in step 3, leading to Figure B2, it becomes clear that the yellow ROI corresponds to observations describing a hole in the tissue, capturing measurements acquired off-tissue and at very low intensity. Besides correctly projecting these measurements to the low-SNR area of the latent space, our workflow also avoids putting these measurements close to any of the genuine biological ‘legs’ in Figure B1b, maintaining the strength-versus-content separation provided by this SiSAL space. This example demonstrates how our workflow, besides exploring biological tissue structure, can also be used to reveal undesirable technical variation or noise variation for removal.

**Fig. B1:**
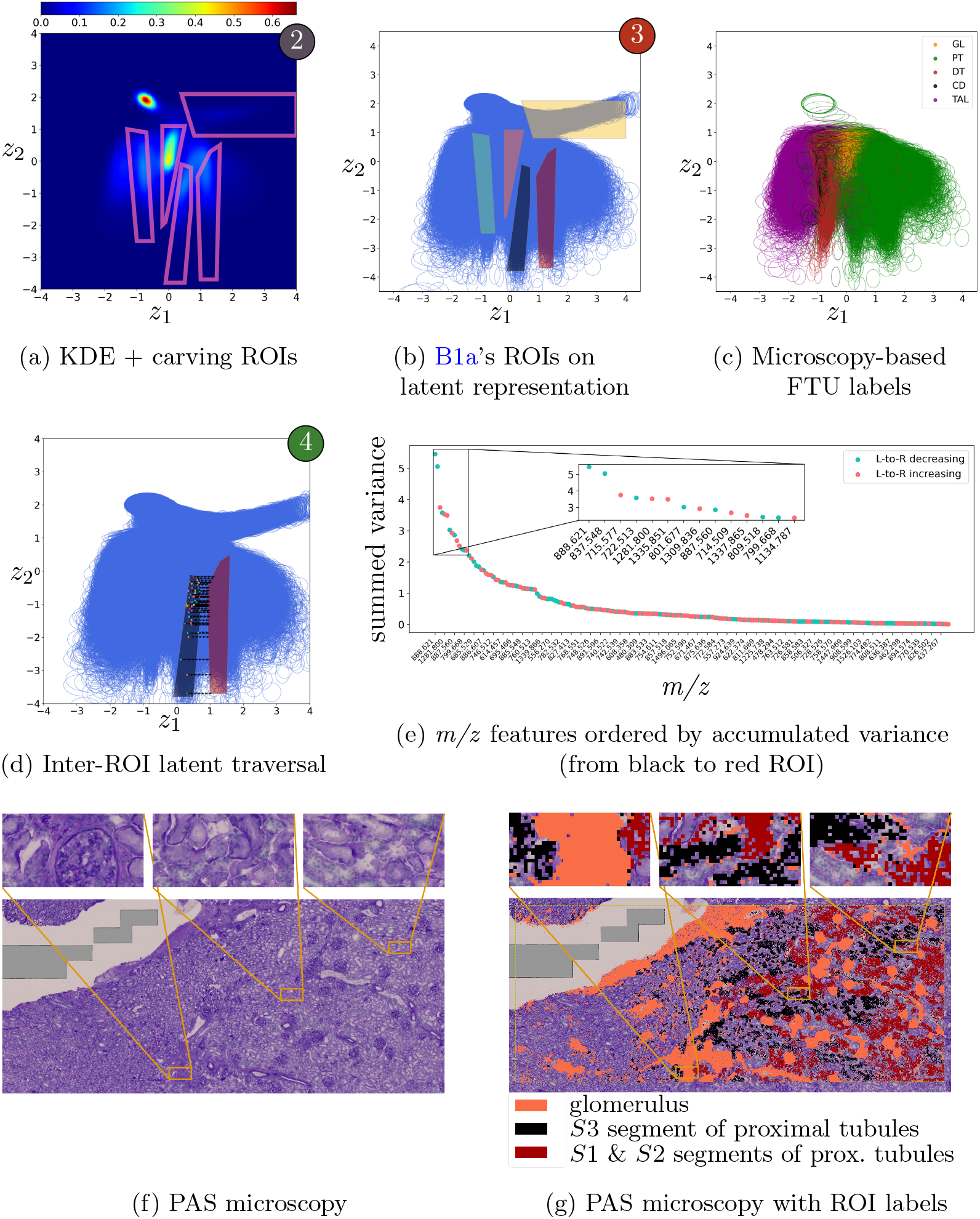
Workflow applied to case study 4’s human kidney data (VAN0042-RK-1-31). Figure B1a, B1b, and B1d report steps 2, 3 and 4. The low-dimensional representation of this dataset demonstrates a SiSAL structure: elongated abundance-reporting striations along one axis and the other axis separating these signatures as chemically distinct. Figure B1c colors the latent space using microscopy-based FTU labels, independently confirming that latent ‘legs’ correspond to different biological structures. The black and red ROIs are defined to examine subdivisions among green PT-reporting pixels. Our comparative-latent-traversal algorithm reveals (B1e) which *m/z* -features vary most between these ROIs. In B1f and B1g, we map our finding to PAS microscopy for biological interpretation.

**Fig. B2:**
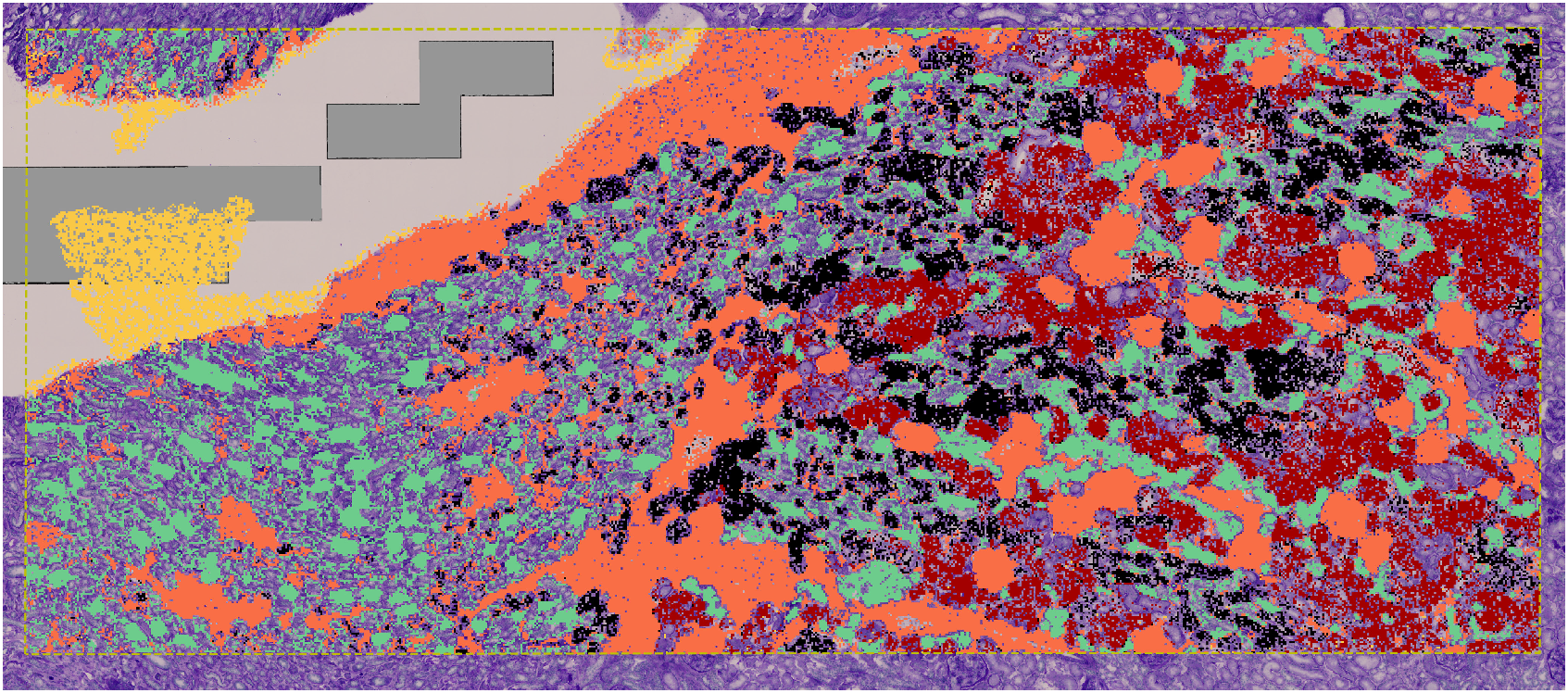
Regions-of-interest from B1b projected onto PAS microscopy

## Supplementary C KDE approximated distribution

### Theorem 1

(Densities change of variables). Let *X* ∈ ℝ*m* be a random variable with density *f*_*X*_ (*z*). Let *µ* : ℝ*m* → ℝ*d* be a mapping with *d* ≤ *m*. Then, *Y* = *µ*(*X*) follows the density 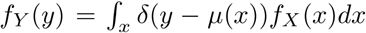, with 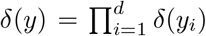, the product of Dirac delta functions.

*Proof*. Let *Z* ∈ ℝ*d* be a random variable with density 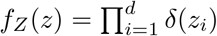, which is a constant vector always equal to 0. Then, we look for the distribution of the random vector *Y* = *µ*(*X*) + *Z*.

The mapping *h* : ℝ*m*+*d* → ℝ*m*+*d* such that *h*(*z, x*) = (*µ*(*x*) + *z, x*) being a bijection, we can define its inverse:

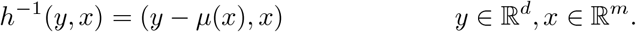

The Jacobian of *h*^−1^ corresponds to

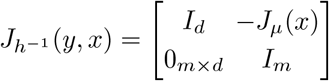

and has a determinant of 1. We thus have:

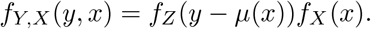

Integrating over *x* gives the desired marginal density of *y*.

## Supplementary D Kullback–Leibler divergence

The following theorem shows a closed form solution for the Kullback-Leibler divergence of two Gaussian distributions. A proof can be found in Section 9 of [29].

### Theorem 2

(Kullback–Leibler divergence of two Gaussian distributions). Let *P* and *Q* be the distributions of two multivariate Gaussian’s 𝒩 (*µ*_1_, Σ_1_) and 𝒩 (*µ*_2_, Σ_2_). We consider *d*-dimensional distributions, meaning that *µ*_1_, *µ*_2_ ∈ ℝ*d* and Σ_1_, Σ_2_ ∈ ℝ*d×d*. The Kullback–Leibler divergence between *P* and *Q* corresponds to:

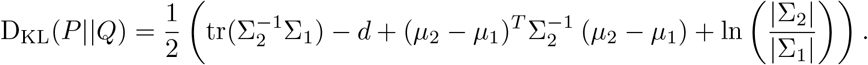

**Fig. B3:**
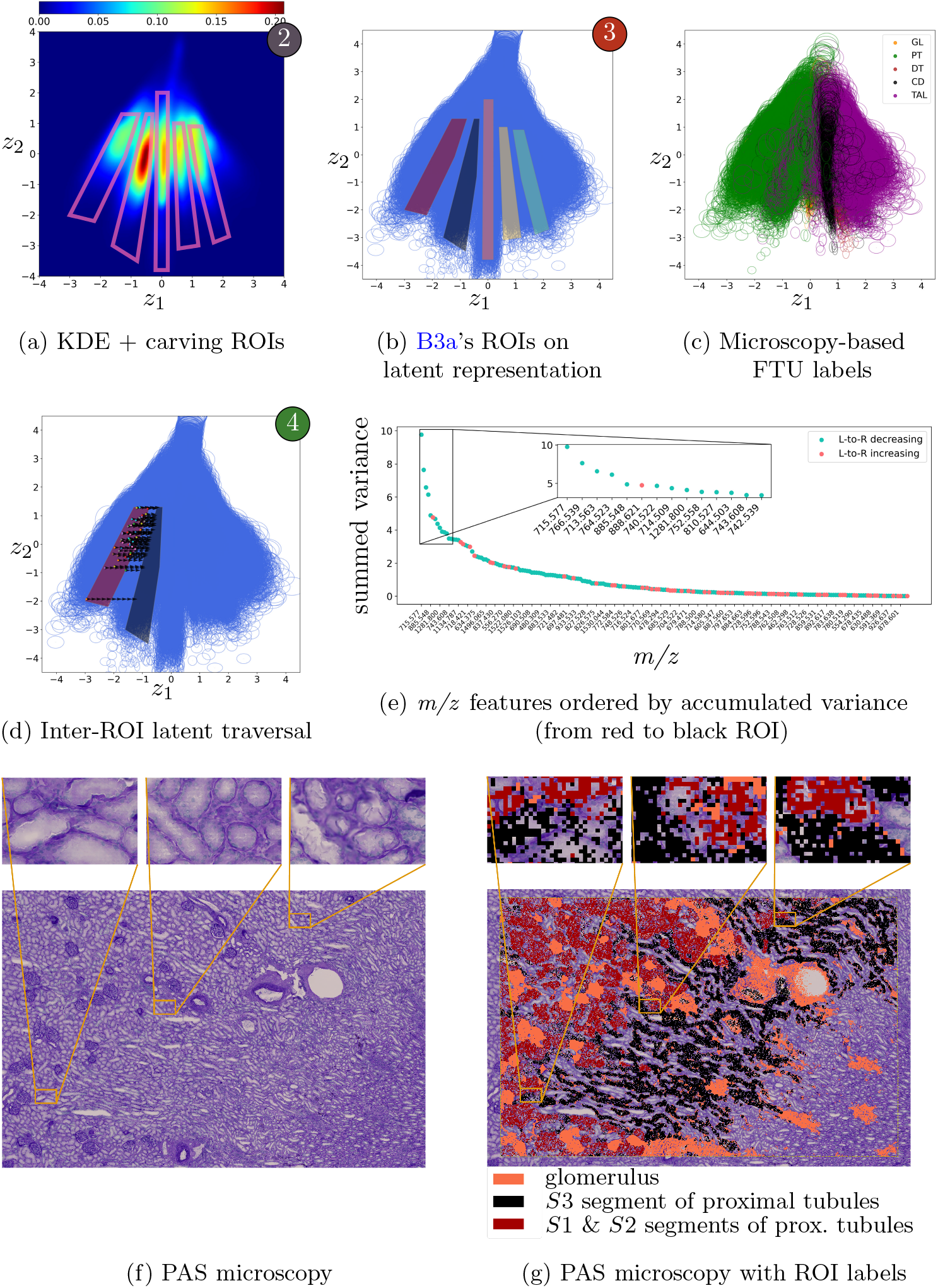
Workflow applied to case study 5’s human kidney data (VAN0049-RK-1-35). Figure B3a, B3b, and B3d report steps 2, 3 and 4. The low-dimensional representation of this dataset demonstrates a SiSAL structure: elongated abundance-reporting striations along one axis and the other axis separating these signatures as chemically distinct. Figure B3c colors the latent space using microscopy-based FTU labels, independently confirming that latent ‘legs’ correspond to different biological structures. The black and red ROIs are defined to examine subdivisions among green PT-reporting pixels. Our comparative-latent-traversal algorithm reveals (B3e) which *m/z* -features vary most between these ROIs. In B3f and B3g, we map our finding to PAS microscopy for biological interpretation.

**Fig. B4:**
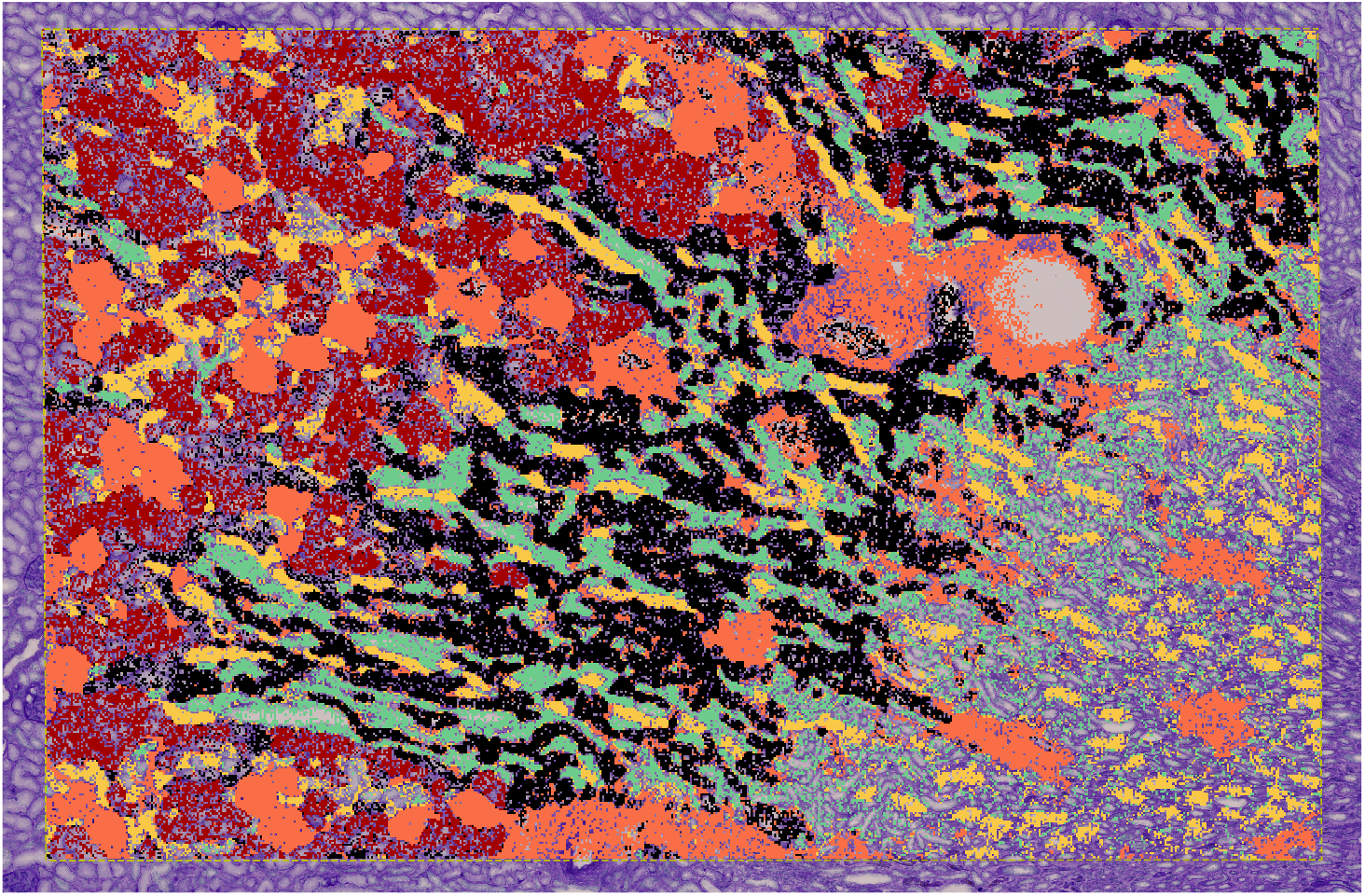
Regions-of-interest from B3b projected onto PAS microscopy

### Theorem 3.

Let *P* a *d*-multivariate Gaussian distribution 𝒩 (*µ*, Σ = Diag *σ*^2^) with *σ* ∈ ℝ*d*, and *Q* be a *d*-standard unitary Gaussian distribution 𝒩 (0, *I*). Then, the *f* (*µ, σ*) = D_KL_(*P* ||*Q*) is a convex function on 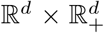, reaching its minimum for *µ*^⋆^ = 0 and *σ*^⋆^ = 1_*d*_.

*Proof*. The Kullback-Leibler divergence between *P* and *Q* corresponds to:

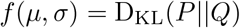

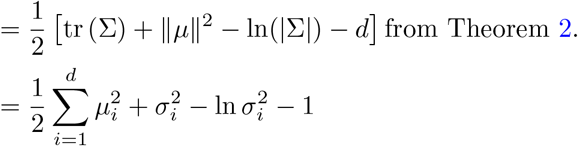

The Hessian matrix of *f* corresponds to:

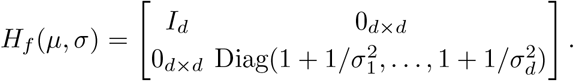

*H*_*f*_ ≻ 0, proving that *f* is convex.

The gradient of *f* is:

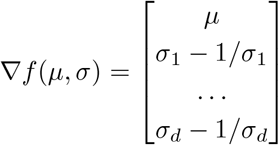

. Setting the gradient to 0 gives us the claimed *µ*^⋆^ and *σ*^⋆^.

## Supplementary E Biological interpretation of molecular differentiators

In the nephron, the PT is exclusively responsible for glucose reabsorption from glomerular filtrate. High capacity, low-affinity SGLT2 is expressed in the early S1+S2 region of the PT, while low-capacity, high-affinity SGLT1 is found in the late S3 PT segment [30]. While both are regulated by the activity of the Na+/K+ ATPase, which establishes the necessary intracellular sodium gradient to drive the uptake of glucose from the lumen, the Na+ and glucose microenvironment of S1+S2 is vastly different from S3. S1+S2 reabsorbs 90% of glucose, and active Na+ transport dominates in this segment. The stoichiometry of SGLT2 is 1:1 Na+ to glucose [30]). The remaining glucose is absorbed in S3, where paracellular Na+ uptake is significantly higher and luminal to basolateral Na+ movement is evenly distributed across passive and active transport mechanisms [31]. Here, the ratio is 2:1 Na+ to glucose. Clearly, the ability of Na+/K+ ATPase to modulate the sodium gradient necessary to drive the influx of glucose is crucial, particularly in the S1+S2 region where SGLT2 activity is high and the Na+ electrochemical gradient is primarily driven by cellular pumps, including Na+/K+ ATPase. Polyunsaturated phosphatidylethonalamine (PE) species have previously been shown to regulate Na+/K+ ATPase activity [26]. Consistent with this, PE (34:2) (*m/z* 714.509 in Fig. 4c, 5g, B1, B3), PE(36:4) (*m/z* 738.508 in Fig. 4c,5g,B1,B3), PE(38:5) (*m/z* 764.523 in Fig. 5g, B3), and PE(38:4) (*m/z* 766.539 in Fig. 5g, B3), were highly correlated with the S1+S2 region. Additionally, PE(38:4) (*m/z* 766.539) has been shown to directly stimulate Na+/K+ ATPase activity [26]. The ability to fine-tune the activity of Na+/K+ ATPase is critical in the regulation of glucose uptake, and the phospholipid environment of Na+/K+ ATPase is a key contributor to pump activity. In addition to Na+/K+ ATPase pump activity, the SGLT transporters themselves are regulated by protein kinases and the phospholipid content of membranes. Studies have shown that protein kinase C (PKC) can upregulate the membrane concentration of SGLT1, but the mechanism for this is unknown, and it is unclear if this is a direct effect of phosphorylation [32]. In contrast, SGLT2 was shown to be phosphorylated by PKC under conditions of insulin stimulation, which induces increased SGLT2 activity [32]. While PKC is recruited to rigid membrane regions where SGLT2 are localized, its activation is inhibited. Alleviation of PKC inhibition is facilitated by phosphatidylserine (PS) binding to PKC [33]. PS(38:4) (*m/z* 810.527 in Fig. 4c, 5g, B3) is exclusively associated with the S1+S2 region where SGLT2 is expressed. Enrichment of this lipid species may be fundamental to PKC activation, leading to SGLT2 phosphorylation and upregulation in S1+S2. Finally, the phospholipid environment itself has been shown to have direct effects on the regulation of SGLT1. Negatively charged lipids were shown to inhibit SGLT1 with phosphatidylinositol (PI) being the most potent inhibitors [34]. However, downregulation of SGLT2 was not observed to be linked to changes in PI lipid content [35]. PI(38:4) (*m/z* 885.548 in Fig 4c, 5g, B3) is localized to S1+S2. This colocalization of PI(38:4) with SGLT2 inhibitors in the S1/S2 segments suggests that PI(38:4) may also influence SGLT2 activity, potentially through the PI3K signaling pathway. Interestingly, PI(38:4) is the most common building block for PIP2 generation [36], and PIP2 has been implicated in endocytosis of the SGLT1 protein, removing it from the plasma membrane [37]. Together, these inhibitory effects of PI on SGLT1 may explain its exclusion from the S3 segment.

## Supplementary F Other dimensionality reduction techniques

In section 2.2, we consider the interpretability of the SiSAL space. For reference, this section shows the results one gets when applying t-SNE (Figure F5) or UMAP (Figure F6) to that same synthetic dataset (Figure 3a). We used well-established implementations of t-SNE [38] and UMAP [39], and we explored a broad variety of hyperparameter settings to avoid drawing parameter-specific conclusions. Both dimensionality reduction techniques seem able to separate dominating components in the data, but their latent axes do not facilitate direct structural interpretation of either the signal content or the SNR.

**Fig. F5:**
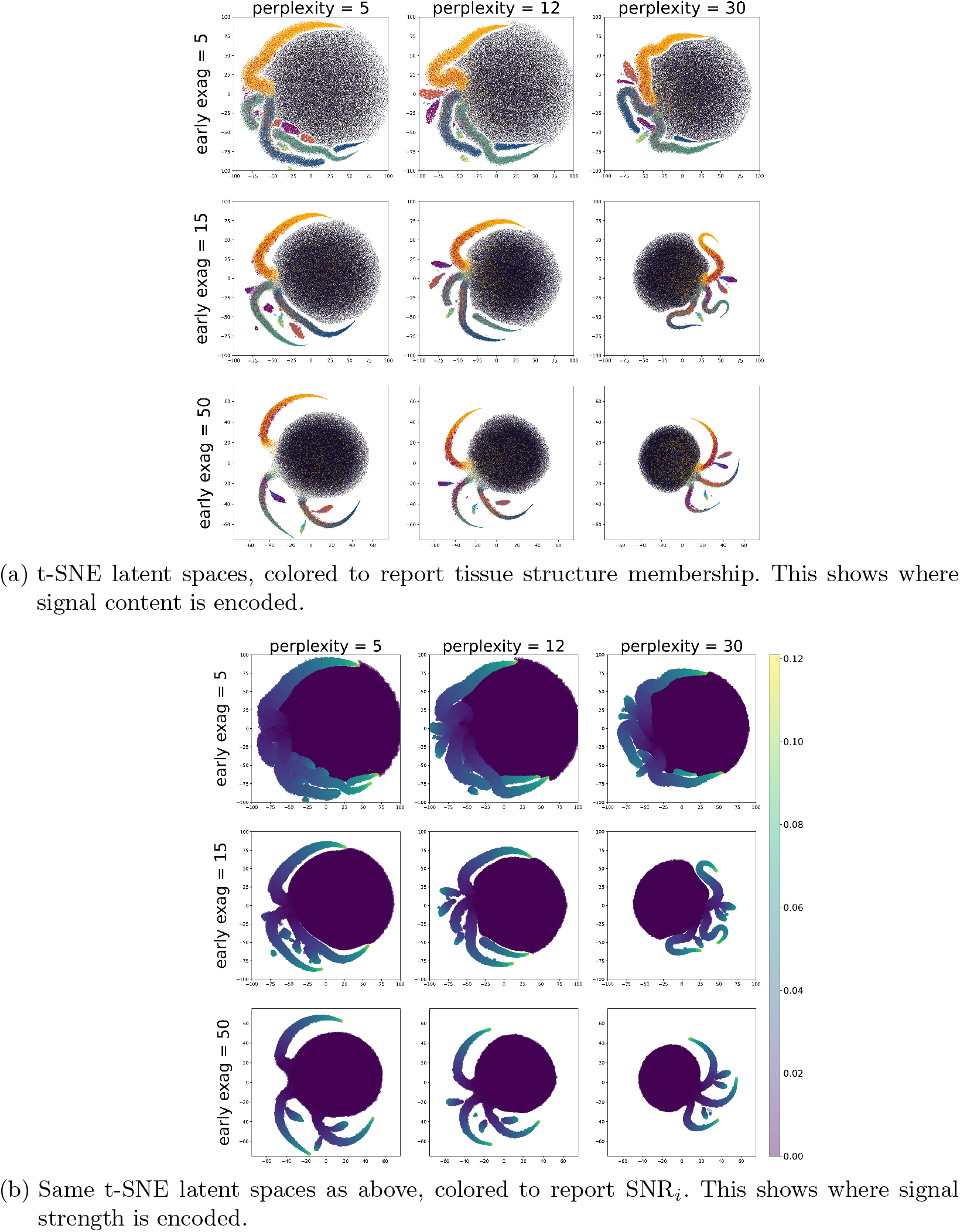
Low-dimensional representations of the synthetic dataset of Figure 3a obtained by t-SNE. Different choices of hyperparameters *perplexity* and *early exageration* are explored to avoid parameter-specific observations. Figures 3b and 3c demonstrate that the SiSAL space captures and encodes signal content and signal strength along distinct latent axes. In the t-SNE-based latent spaces, such separation along latent axes is not observed, demonstrating the structural nature and interpretability of the SiSAL space compared to traditional methods.

**Fig. F6:**
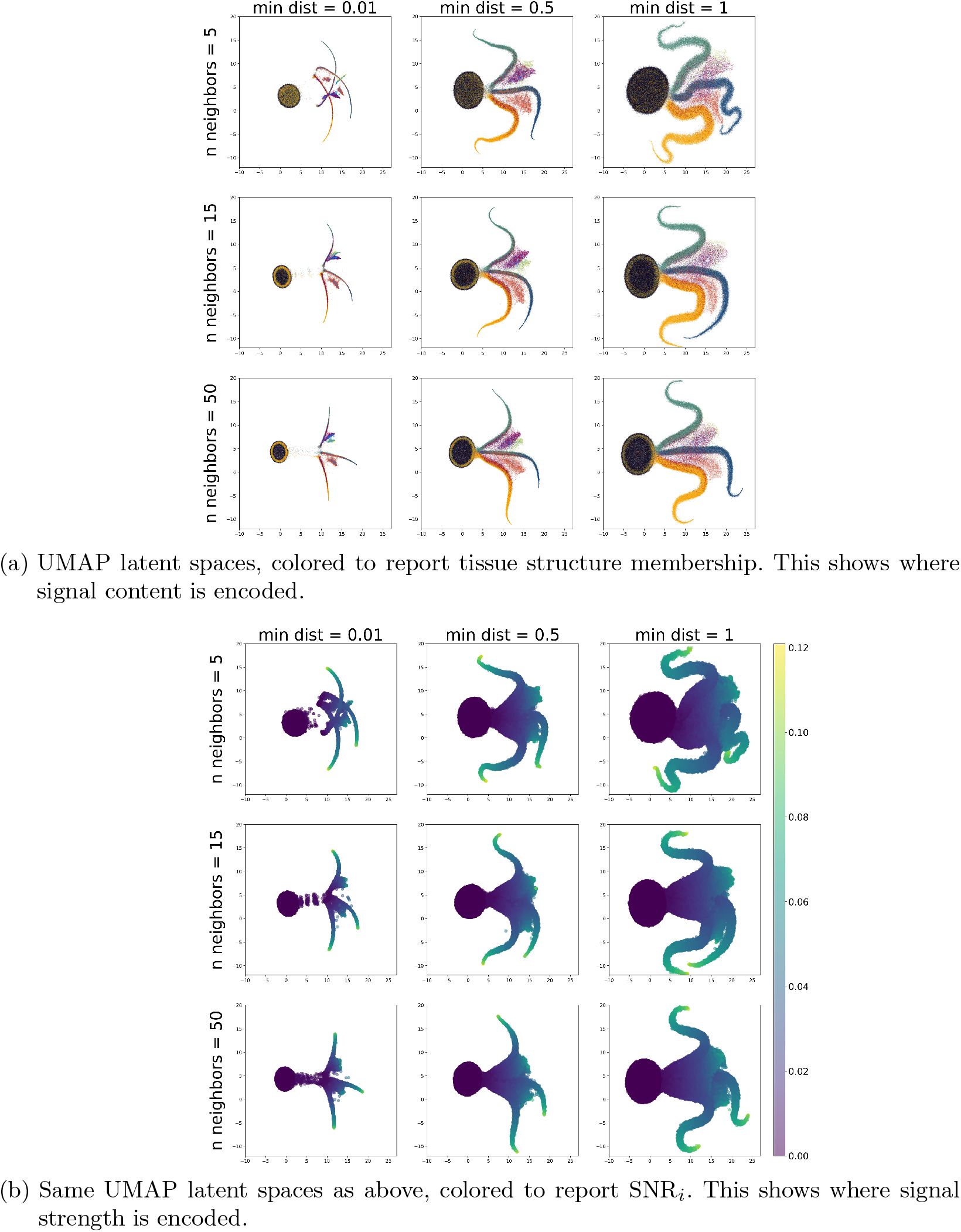
Low-dimensional representations of the synthetic dataset of Figure 3a obtained by UMAP. Different choices of hyperparameters *number of neighbors* and *minimum distance* are explored to avoid parameter-specific observations. Figures 3b and 3c demonstrate that the SiSAL space captures and encodes signal content and signal strength along distinct latent axes. In the UMAP-based latent spaces, such separation along latent axes is not observed, demonstrating the structural nature and interpretability of the SiSAL space compared to traditional methods.

## References

[1] Caprioli, R.M., Farmer, T.B., Gile, J.: Molecular imaging of biological samples: localization of peptides and proteins using maldi-tof ms. Analytical chemistry 69(23), 4751–4760 (1997)

[2] McDonnell, L.A., Heeren, R.M.: Imaging mass spectrometry. Mass spectrometry reviews 26(4), 606–643 (2007)

[3] Ren, J., Zabalza, J., Marshall, S., Zheng, J.: Effective feature extraction and data reduction in remote sensing using hyperspectral imaging [applications corner]. IEEE Signal Processing Magazine 31(4), 149–154 (2014)

[4] Deng, X., Tian, X., Chen, S., Harris, C.J.: Deep principal component analysis based on layerwise feature extraction and its application to nonlinear process monitoring. IEEE Transactions on Control Systems Technology 27(6), 2526–2540 (2018)

[5] Kharchenko, P.V.: The triumphs and limitations of computational methods for scrna-seq. Nature methods 18(7), 723–732 (2021)

[6] Weylandt, M., Swiler, L.P.: Beyond pca: Additional dimension reduction techniques to consider in the development of climate fingerprints. Journal of Climate 37(5), 1723–1735 (2024)

[7] Pearson, K.: Liii. on lines and planes of closest fit to systems of points in space. The London, Edinburgh, and Dublin philosophical magazine and journal of science 2(11), 559–572 (1901)

[8] Hotelling, H.: Analysis of a complex of statistical variables into principal components. Journal of educational psychology 24(6), 417 (1933)

[9] Principal Component Analysis for Special Types of Data, pp. 338–372. Springer, New York, NY (2002). 10.1007/0-387-22440-8_13. https://doi.org/10.1007/0-387-22440-8_13

[10] Palmer, A.D., Bunch, J., Styles, I.B.: Randomized approximation methods for the efficient compression and analysis of hyperspectral data. Anal Chem 85(10), 5078–86 (2013)

[11] Lee, D.D., Seung, H.S.: Learning the parts of objects by non-negative matrix factorization. Nature 401(6755), 788–791 (1999)

[12] Lee, D., Seung, H.S.: Algorithms for non-negative matrix factorization. Advances in neural information processing systems 13 (2000)

[13] Maaten, L., Hinton, G.: Visualizing data using t-sne. Journal of Machine Learning Research 9(86), 2579–2605 (2008)

[14] McInnes, L., Healy, J., Melville, J.: UMAP: Uniform Manifold Approximation and Projection for Dimension Reduction (2020)

[15] Clarke, H.A., Ma, X., Shedlock, C.J., Medina, T., Hawkinson, T.R., Wu, L., Ribas, R.A., Keohane, S., Ravi, S., Bizon, J.L., Burke, S.N., Abisambra, J.F., Merritt, M.E., Prentice, B.M., Vander Kooi, C.W., Gentry, M.S., Chen, L., Sun, R.C.: Spatial mapping of the brain metabolome lipidome and glycome. Nat Commun 16(1), 4373 (2025)

[16] Kingma, D.P., Welling, M.: Auto-encoding variational bayes. stat 1050, 1 (2014)

[17] Higgins, I., Matthey, L., Pal, A., Burgess, C., Glorot, X., Botvinick, M., Mohamed, S., Lerchner, A.: beta-vae: Learning basic visual concepts with a constrained variational framework. In: International Conference on Learning Representations (2016)

[18] Thomas, S.A., Race, A.M., Steven, R.T., Gilmore, I.S., Bunch, J.: Dimensionality reduction of mass spectrometry imaging data using autoencoders. In: 2016 IEEE Symposium Series on Computational Intelligence (SSCI), pp. 1–7 (2016). 10.1109/SSCI.2016.7849863

[19] Guo, D., Föll, M.C., Bemis, K.A., Vitek, O.: A noise-robust deep clustering of biomolecular ions improves interpretability of mass spectrometric images. Bioinformatics 39(2), 067 (2023) 10.1093/bioinformatics/btad067 https://academic.oup.com/bioinformatics/article-pdf/39/2/btad067/49286651/btad067.pdf

[20] Abdelmoula, W.M., Lopez, B.G.-C., Randall, E.C., Kapur, T., Sarkaria, J.N., White, F.M., Agar, J.N., Wells, W.M., Agar, N.Y.: Peak learning of mass spectrometry imaging data using artificial neural networks. Nature communications 12(1), 5544 (2021)

[21] Inglese, P., Alexander, J.L., Mroz, A., Takats, Z., Glen, R.: Variational autoencoders for tissue heterogeneity exploration from (almost) no preprocessed mass spectrometry imaging data. arXiv preprint 1708.07012 (2017)

[22] Jain, S., Pei, L., Spraggins, J.M., Angelo, M., Carson, J.P., Gehlenborg, N., Ginty, F., Gonçalves, J.P., Hagood, J.S., Hickey, J.W., et al.: Advances and prospects for the human biomolecular atlas program (hubmap). Nature cell biology, 1–12 (2023)

[23] HuBMAP Consortium: The human body at cellular resolution: the nih human biomolecular atlas program. Nature 574(7777), 187–192 (2019)

[24] Delacour, P.-L., Wahls, S., Spraggins, J.M., Migas, L., Van-de-Plas, R.: Signal recovery using a spiked mixture model. IEEE Transactions on Signal Processing, 1–14 (2025) 10.1109/TSP.2025.3593082

[25] Patterson, N.H., Neumann, E.K., Sharman, K., Allen, J., Harris, R., Fogo, A.B., Caestecker, M., Caprioli, R.M., Plas, R., Spraggins, J.M.: Autofluorescence microscopy as a label-free tool for renal histology and glomerular segmentation. bioRxiv, 2021–07 (2021)

[26] Habeck, M., Haviv, H., Katz, A., Kapri-Pardes, E., Ayciriex, S., Shevchenko, A., Ogawa, H., Toyoshima, C., Karlish, S.J.: Stimulation, inhibition, or stabilization of na, k-atpase caused by specific lipid interactions at distinct sites. Journal of Biological Chemistry 290(8), 4829–4842 (2015) 10.1074/jbc.M114.611384

[27] Goodfellow, I., Bengio, Y., Courville, A.: Deep Learning, (2016).Chap. 19

[28] Scott, D.W.: Multivariate density estimation: theory, practice, and visualization. John Wiley & Sons (2015)

[29] Duchi, J.: Derivations for linear algebra and optimization

[30] Wright, E.M., Loo, D.D.F., Hirayama, B.A.: Biology of human sodium glucose transporters. Physiological Reviews 91, 733–794 (2011) 10.1152/physrev.00055.2009

[31] Layton, A.T., Vallon, V., Edwards, A.: Modeling oxygen consumption in the proximal tubule: effects of nhe and sglt2 inhibition. American Journal of Physiology-Renal Physiology 308, 1343–1357 (2015) 10.1152/ajprenal.00007.2015

[32] Ghezzi, C., Wright, E.M.: Regulation of the human na+-dependent glucose cotransporter hsglt2. American Journal of Physiology-Cell Physiology 303, 348–354 (2012) 10.1152/ajpcell.00115.2012

[33] Jiang, Y., Pan, Z., Chen, J.W.: Interaction between protein kinase c and sphingomyelin/cholesterol. Cell Biology International 23(7), 457–463 (1999) 10.1006/cbir.1999.0374

[34] Ebel, H., Fromm, A., Günzel, D., Fromm, M., Schulzke, J.D.: Phospholipid effects on sglt1-mediated glucose transport in rabbit ileum brush border membrane vesicles. Biochimica et Biophysica Acta (BBA) Biomembranes 1861(10), 182985 (2019) 10.1016/j.bbamem.2019.05.007

[35] Albertoni Borghese, M.F., Majowicz, M.P., Ortiz, M.C., Rosario Passalacqua, M., Sterin Speziale, N.B., Vidal, N.A.: Expression and activity of sglt2 in diabetes induced by streptozotocin: Relationship with the lipid environment. Nephron Physiology 112(3), 45–52 (2009) 10.1159/000214214 https://karger.com/nep/article-pdf/112/3/p45/3776683/000214214.pdf

[36] Borges-Araújo, L., Domingues, M.M., Fedorov, A., Santos, N.C., Melo, M.N., Fernandes, F.: Acyl-chain saturation regulates the order of phosphatidylinositol 4, 5-bisphosphate nanodomains. Communications Chemistry 4(1), 164 (2021) 10.1038/s42004-021-00603-1

[37] Ghezzi, C., Calmettes, G., Morand, P., Ribalet, B., John, S.: Real-time imaging of sodium glucose transporter (sglt1) trafficking and activity in single cells. Physiological reports 5(3), 13062 (2017) 10.14814/phy2.13062

[38] Pedregosa, F., Varoquaux, G., Gramfort, A., Michel, V., Thirion, B., Grisel, O., Blondel, M., Prettenhofer, P., Weiss, R., Dubourg, V., Vanderplas, J., Passos, A., Cournapeau, D., Brucher, M., Perrot, M., Duchesnay, E.: Scikit-learn: Machine learning in Python. Journal of Machine Learning Research 12, 2825–2830 (2011)

[39] McInnes, L., Healy, J., Saul, N., Grossberger, L.: Umap: Uniform manifold approximation and projection. The Journal of Open Source Software 3(29), 861 (2018)

